# Human Telomerase Expression is under Direct Transcriptional Control of the Telomere-binding-factor TRF2

**DOI:** 10.1101/2020.01.15.907626

**Authors:** Shalu Sharma, Ananda Kishore Mukherjee, Shuvra Shekhar Roy, Sulochana Bagri, Silje Lier, Meenakshi Verma, Antara Sengupta, Manish Kumar, Gaute Nesse, Deo Prakash Pandey, Shantanu Chowdhury

**Affiliations:** Integrative and Functional Biology Unit, CSIR- Institute of Genomics and Integrative Biology, Mathura Road, New Delhi-110025, India; Academy of Scientific and Innovative Research, CSIR- Institute of Genomics and Integrative Biology, Mathura Road, New Delhi-110025, India; GNR Knowledge Centre for Genome and Informatics, CSIR- Institute of Genomics and Integrative Biology, Mathura Road, New Delhi-110025, India; Imaging Facility CSIR-IGIB, CSIR- Institute of Genomics and Integrative Biology, Mathura Road, New Delhi-110025, India; Dept. of Microbiology, Oslo University Hospital, Oslo, Norway, CSIR- Institute of Genomics and Integrative Biology, Mathura Road, New Delhi-110025, India; Institute of basic medical sciences, University of Oslo, Oslo, CSIR- Institute of Genomics and Integrative Biology, Mathura Road, New Delhi-110025, India

## Abstract

Tight regulatory mechanisms to maintain repression of human Telomerase (hTERT), the sole protein that synthesizes telomeres, is crucial for normal adult somatic cells. In contrast, enhanced telomerase activity and resulting pathological maintenance of telomeres, is widely understood as causal in >90% of human cancers. These implicate underlying mechanisms connecting *hTERT* regulation and telomeres, possibly through telomeric proteins, that remain unclear. In light of of recent work by us and others showing non-telomeric function of the telomere-binding protein TRF2, here we examined whether and how TRF2 affected *hTERT* regulation. Direct binding of TRF2 – spanning ∼450 bp of the *hTERT* promoter from the Transcriptional Start Site (TSS) – led to TRF2-dependent recruitment of the polycomb repressor complex PRC2 in both normal and cancer cells. This induced repressor histone modifications resulting in TRF2-dependent *hTERT* repression. Mutations in the *hTERT* promoter, found frequently in aggressive glioblastoma and reported to destabilize the G-quadruplex structure, resulted in loss of TRF2 binding and consequent *hTERT* over-expression. Conversely, using G-quadruplex-stabilizing ligands we regained TRF2 binding, *hTERT* re-suppression, in highly proliferating glioblastoma cells with telomerase hyperactivation due to *hTERT* promoter mutations. Together, results herein demonstrate direct control of *hTERT* through TRF2 in a G-quadruplex-dependent manner – implicating mechanisms of how telomerase regulation might be linked to telomeres in normal and cancer cells.

## Introduction

Telomeres, comprising of (GGGTTA)_n_ repeats in complex with telomere-binding proteins, at the end of human chromosomes are essential for genome stability ^1–6^. The only protein that can replicate telomeric DNA is telomerase - a ribonucleoprotein complex of the reverse transcriptase (hTERT) and RNA the template (hTERC). In adult somatic cells, *hTERT* is kept trasncriptionally repressed. Resulting loss of telomeres with each replication cycle leads to replicative senescence, much like a ‘molecular clock’ that helps maintain cellular homeostasis. Deregulation or loss of *hTERT* repression, which results in aberrant maintenance of telomeres, has been causally associated to initiation/progression of more than 90% human cancers^7,8^. Although, these suggest tight control of telomerase might be linked to telomeres - the role of telomeres or telomere-binding factors in regulation of *hTERT* remains poorly explored.

Relatively recent work by others and us showing non-telomeric binding of telomere-binding proteins, TRF1, TRF2 and RAP1, is of interest in this context. Genome wide RAP1 association has been reported in mouse and human cells while TRF1/TRF2 binding has been demonstrated in human cells ^9–11^. Notably, we found about 20000 TRF2 binding sites spread throughout the genome where TRF2-mediated promoter epigenetics and gene regulation was evident. A large fraction of the TRF2 binding sites coincided with potential DNA secondary structure G-quadruplex-forming sequences^12^. Further, we observed interaction of TRF2 with G-quadruplexes at multiple promoters, consistent with the emerging role of G-quadruplexes as epigenetic regulatory motifs (reviewed in^13–17^). Moreover, the *hTERT* near promoter was reported to harbor multiple G-quadruplexes^18–20^.

Here, we asked if TRF2 directly associates with the *hTERT* promoter and affects regulation of *hTERT* expression. We found direct binding of TRF2 at the *hTERT* promoter and TRF2-mediated transcriptional repression of *hTERT*. This was evident from experiments focussed on the endogenous *hTERT* in multiple normal and cancer cell types, as well as an exogenously introduced *hTERT* promoter-reporter cassette (CRISPR-Cas-mediated insertion). Furthermore, recruitment of the epigenetic polycomb repressor complex PRC2 at the *hTERT* promoter was observed to be TRF2-dependent, and necessary for retaining chromatin in a non-permissive state at the *hTERT* loci. Importantly, TRF2 promoter binding was dependent on the presence of intact G-quadruplex. Notably, in glioblastoma (GBM) patient-derived cells with *hTERT* promoter mutation(s), that destabilize G-quadruplex(es)^21^ – reported to be frequently associated with aggressive GBM and other human cancers^22^ - G-quadruplex-binding ligands regained promoter TRF2 occupancy, reinstated the epigenetic repressor machinery and resulted in *hTERT* repression in a TRF2-dependent fashion. Together, these demonstrate a new link between telomeres and telomerase expression through non-telomeric binding of TRF2. The resulting repressive chromatin at the *hTERT* promoter might be crucial for keeping normal human somatic cells from re-activating telomerase and triggering cancer in cells.

## Results

### TRF2 directly associates with the *hTERT* promoter and suppresses *hTERT* expression

We recently reported TRF2 ChIP-seq peaks throughout the genome^13^. Interestingly, TRF2 ChIP-seq reads were noted within the *hTERT* near promoter (Supplementary Fig 1A). Here we tested TRF2 occupancy on the *hTERT* promoter in human cancer (fibrosarcoma HT1080 and colon carcinoma HCT116 cells) and normal (primary fibroblast MRC5 and human embryonic kidney HEK293T) cells directly. ChIP-PCR showed TRF2 binding on the *hTERT* promoter spanning a region from the transcription start site (TSS) to 750 bp upstream in all four cell lines (Figure 1A).

**Figure 1.**
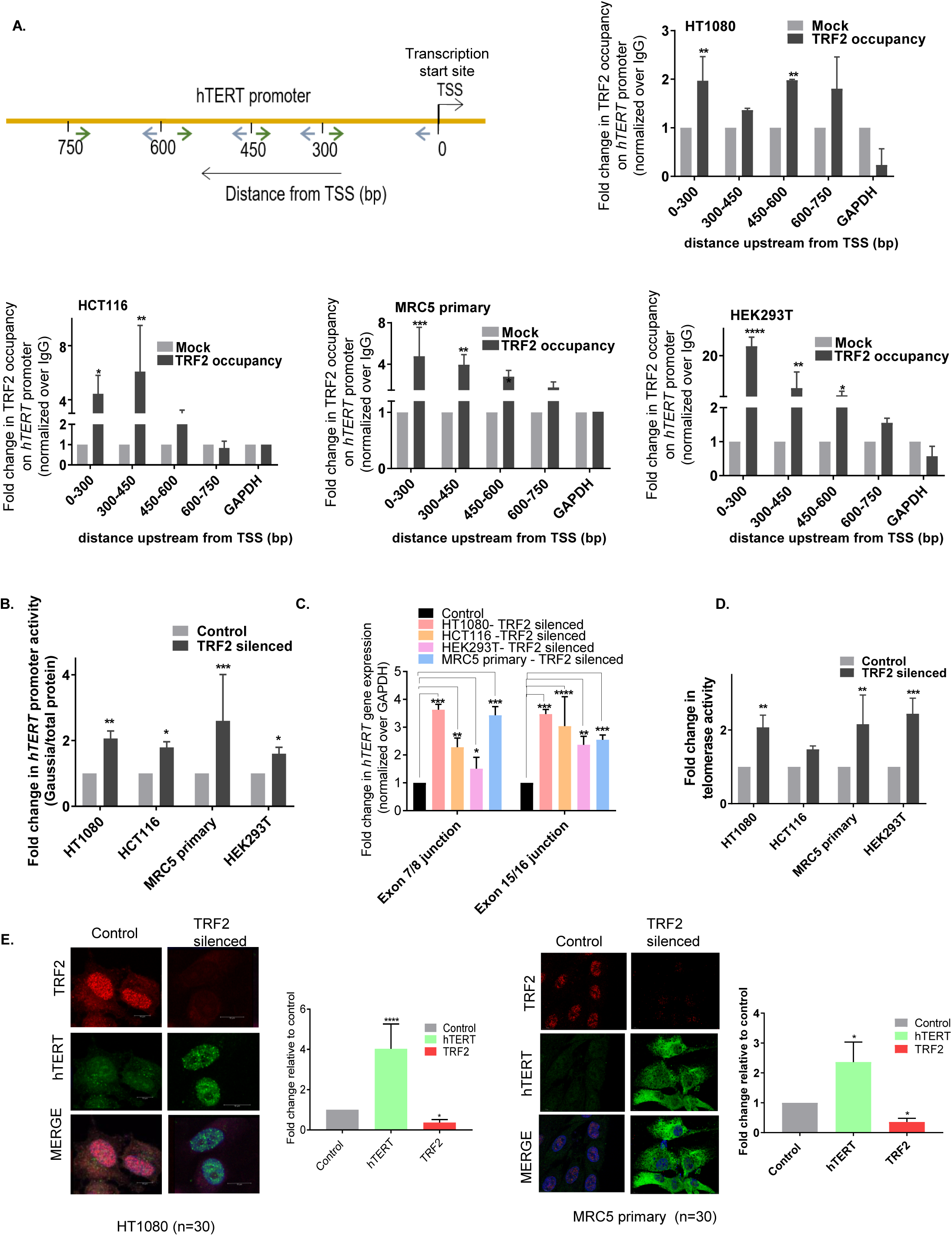
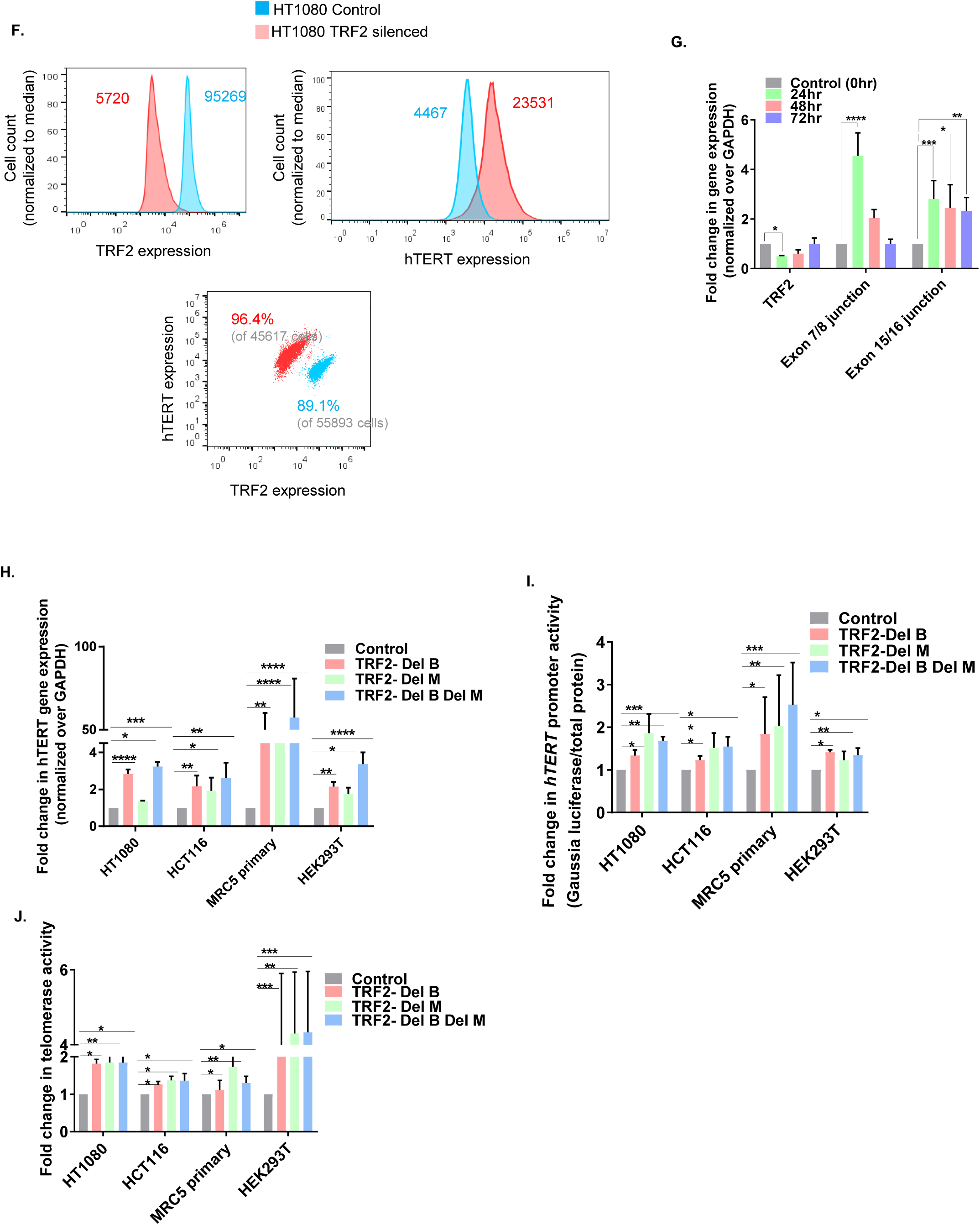
TRF2 binds at the *hTERT* promoter and this results in transcriptionally repression of *hTERT*. **A**. *hTERT* promoter, spanning primers from 0 to -750bp upstream from the transcription start site (TSS); TRF2 ChIP followed by *hTERT*-promoter spanning qRT-PCR, showing TRF2 binding in HT1080 and HCT116 cancer cells, primary fibroblast MRC5 cells and transformed normal HEK293T cells. **B**. Effect of siRNA mediated TRF2 silencing on *hTERT* promoter activity, +33 to-1276 bp *hTERT* promoter cloned upstream of Gaussia luciferase in HT1080, HCT116, MRC5 primary and HEK293T cells post 48hrs of transfection. **C**. Effect of TRF2 silencing on *hTERT* expression (functional transcript-exon 7-8 and exon 15-16 full transcript) in HT1080, HCT116, MRC5 and HEK293T cells. **D**. Effect of TRF2 silencing on telomerase activity in HT1080, HCT116, MRC5 primary and HEK293T cells measure using ELISA TRAP (see methods) signal ws normalized over untreated cells (control). **E**. Immunofluorescence staining of hTERT and TRF2 protein in HT1080 and MRC5 primary cells (N=30) TRF2 and hTERT were stained using Alexa fluor594 (red signal) and Alexa fluor 498 (green signal) respectively, quantification of signal from N=30 cells is shown in the right panel, normalized over control (untreated cells). **F**. Flow cytometric analysis using dual staining of hTERT and TRF2 in HT1080 control cells in comparison to TRF2 silenced cells, gain in Mean intensity of Fluroscence (MIF) of hTERT was measured with loss in MIF of TRF2 upon TRF2 silencing, below 89.1% of 55893 cells taken from control population showed higher TRF2 and lower *hTERT* the trend reversed on TRF2 silencing in 96.4% population of 45617 cells. **G**. Expression of *TRF2* and *hTERT* gene (exon7-8 and exon 15-16) was tested at 24, 48 and 72hrs following TRF2 siRNA treatment; the siRNA complex was removed 6hrs post transfection. **H-J**. Expression of hTERT transcript (exon 15-16) **H**.; *hTERT* promoter activity **I**. and telomerase activity **J**. following expression of TRF2 deletion mutants lacking DNA binding activity: each case results were normalized to untreated control cells. All error bars represent ± standard deviations from mean values and p values have been calculated by paired /un paired T-test. (*: p<0.05, **: p<0.01, ***: p<0.005, ****: p<0.0001).

Next, we checked whether TRF2 altered transcription of *hTERT*. Promoter activity (from +33 to -1267 bp *hTERT* promoter introduced upstream of Gaussia luciferase reporter construct) was markedly enhancedupon siRNA-mediated TRF2 silencing in all the four cell types (Figure 1B). Consistent with this, endogenous *hTERT* expression was upregulated in both normal and cancer cells on TRF2 silencing; expression of both, the functional reverse transcriptase domain (exon 7/8) as well as the full-length *hTERT* transcript (exon 15/16) was enhanced (Figure 1 C). For further confirmation, we tested the effect of TRF2 on intracellular telomerase activity. Silencing of TRF2 resulted in enhanced telomerase activity in all the four cell lines (Figure 1D); and, increase in telomerase protein on TRF2 silencing was also evident (Supplementary Figure 1B).

Next, we performed immunofluorescence (IF) experiments to check hTERT protein levels within single cells in HT1080 and primary MRC5 cells. On TRF2 silencing two-three fold enhanced hTERT levels was evident in both cell types (Figure 1E). In HT1080 cancer cells, enhanced telomerase was found mostly within the nuclei as expected; however, in case of the primary MRC5 cells this was observed outside of the nuclei also, as noted earlier^23,24^.

For further validation flow-cytometry analysis (fluorescence-assisted cell sorting (FACS)) of hTERT levels following silencing of TRF2 (using TRF2 siRNA as above) in HT1080 cells was done. The mean fluorescence intensity (MFI) of TRF2 decreased by ∼16 folds, whereas the hTERT-MFI increased by ∼5.2 fold, in cells where TRF2 was silenced relative to untreated control cells (Figure 1F). The cell populations monitored were: control (89.1% of 55893) and TRF2-silenced (96.4% of 45716) where staining of TRF2-high: hTERT-low and TRF2-low: hTERT-high cells, respectively, was clear (Figure 1F).

In addition to this, we performed rescue experiments using TRF2 siRNA. TRF2 was first depleted in HT1080 cells using siRNA, which gave enhanced *hTERT* expression. Thereafter, cells were maintained for 72 hrs with no further siRNA addition when TRF2 levels gradually increased - concomitant decline in *hTERT* was clearly evident (Figure 1G). Taken together, these suggested transcriptional control of functional telomerase by TRF2. The antibody used here for hTERT was confirmed by FACS in super-telomerase cells that constitutively over-express telomerase (characterized earlier^25^) (Supplementary Figure 1C).

### DNA binding by TRF2 is necessary for transcription regulation of *hTERT*

We next asked if direct DNA binding by TRF2 is necessary for TRF2-mediated repression of *hTERT*. To test this, we over-expressed Flag-tagged TRF2-DelM (lacking C-terminal Myb (M) domain), TRF2-DelB (lacking N-terminal Basic (B) domain) or TRF2-DelB-DelM (lacking both B and M domains domains) mutants of TRF2 that lack DNA binding. In all the three mutants we observed enhanced expression of endogenous *hTERT* full transcript in HT1080, HCT116 and MRC5 primary and HEK293T cells (Figure 1H). In addition, over-expression of TRF2-DelM, TRF2-DelB and TRF2-DelB-DelM also resulted in enhanced promoter luciferase activity of *hTERT* (Figure 1I) and increase in telomerase activity in all the four cell lines (Figure 1J). As expected, binding of the TRF2-DelB-DelM mutant on the *hTERT* promoter was not significant (Supplementary Figure 1D). We also checked, whether the dominant negative effect of TRF2-DelM noted earlier^12^ which was likely to reduce the binding of endogenous full-length TRF2 was observed in case of the *hTERT* promoter. As expected TRF2 occupancy was lost at the *hTERT* promoter on over-expression of TRF2-DelB-DelM (Supplementary, Figure 1D).

### Epigenetic state of chromatin at the *hTERT* promoter is TRF2-dependent

TRF2-mediated change in promoter histone methylation at several promoters spread across the genome was observed earlier^13,26–30^ Here, we sought to understand whether TRF2-mediated transcriptional repression of *hTERT* involved altered epigenetic state at the promoter. Changes in two histone-activation (mono and tri-methylated histone H3 lysine 4 (H3K4me1 and H3K4me3)) and two repressors (tri-methylated histone H3 lysine 9 and lysine 27 (H3K27me3 and H3K9me3)) marks were checked at the *hTERT* promoter following TRF2 silencing. ChIP-PCR for each of the four histone marks using primers spanning a region up to 750 bp upstream of *hTERT* TSS (as in Figure 1A) was performed. Interestingly, we found significant loss in only the H3K27me3 repressor mark in both HT1080 and MRC5 primary cells (Figure 2A and B) whereas activation marks H3K4me1, H3K4me3 or the repressor mark H3K9me3 did not change significantly on TRF2 silencing (Supplementary Figure 2 A and B).

**Figure 2.**
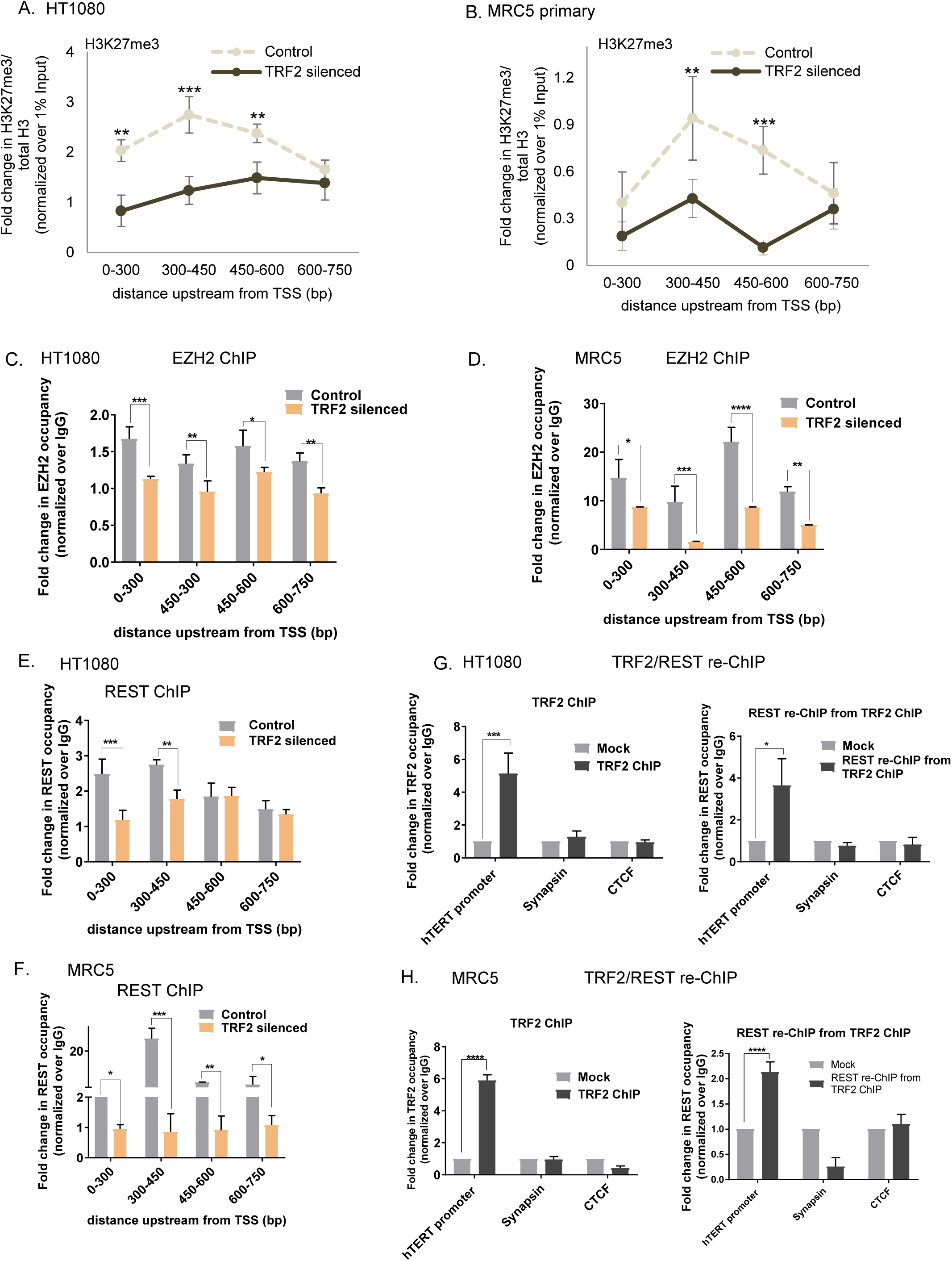
TRF2 recruits the polycomb repressor complex PRC2 at the *hTERT* promoter. **A-B**. Effect of TRF2 silencing on H3K27me3 occupancy spanning 0-750bp on *hTERT* promoter; fold change in H3K27me3 over total H3 normalized over 1% Input **A**. in HT1080 cells, **B**. MRC5 primary cells. **C-D**. Fold change in EZH2 occupancy spanning *hTERT* promoter **C**. HT1080 cells **D**. MRC5 primary cells **E-F**. Fold change in REST occupancy on *hTERT* promoter in **E**. HT1080 cells **F**. MRC5 primary cells. **G-H**. TRF2 ChIP followed by REST re-ChIP: TRF2 ChIP (left panel) and REST re-ChIP (right panel) **G**. in HT1080 and **H**. MRC5 primary cells (*hTERT* core promoter (+38 to -237bp), *Syanapsin* and *CTCF-*negative control for TRF2 binding All error bars represent ± standard deviations from mean values and p values have been calculated by paired /un paired T-test. (*: p<0.05, **: p<0.01, ***: p<0.005, ****: p<0.0001).

### TRF2-mediated recruitment of the polycomb repressor complex PRC2 at the *hTERT* promoter

The polycomb-repressor complex 2 (PRC2) is known to catalyse H3K27me3 modification exclusively resulting in gene inactivation^31–33^. Therefore, we tested if the occupancy of the PRC2 complex on the *hTERT* promoter was TRF2-dependent. In both HT1080 and MRC5 cells we found that TRF2 silencing resulted in loss of EZH2 (catalytic component of the PRC2 complex) occupancy (Figure 2C and D). However, on silencing of EZH2, TRF2 occupancy remained relatively un-altered (Supplementary Figure 2C) suggesting TRF2-dependent recruitment of EZH2/PRC2 complex at the *hTERT* promoter.

Recruitment of the PRC2 complex through the RE1-silencing factor (REST) throughout the genome was observed earlier^34–36^. On the other hand, interaction of TRF2 with REST was observed by us and others^12,26,37–39^. We sought to check, therefore, if TRF2 recruited REST on the *hTERT* promoter. In both HT1080 and MRC5 cells, TRF2 silencing resulted in loss of REST association from the *hTERT* promoter region (up to 750 bp upstream) suggesting TRF2-dependent REST occupancy (Figure 2E, F). This is supported by intracellular interaction of TRF2 and REST noted by us in HT1080 cells earlier^30^ and in MRC5 primary cells shown here through co-immunoprecipitation (co-IP) of REST using anti-TRF2 antibody (Supplementary Figure 2D).

To further substantiate TRF2-dependent REST binding on the *hTERT* promoter we performed re-ChIP experiments: REST-reChIP from the TRF2 ChIP fraction (see methods) in both HT1080 and MRC5 cells showed TRF2-dependent REST occupancy. We used the *CTCF* promoter as a negative control for TRF2 ChIP and the *synapsin* promoter (reported for REST occupancy^40,41^, but not TRF2) as a positive control for the REST ChIP (Figure 2G and 2H). As expected, the REST reChIP was negative for *synapsin*.

Conversely, TRF2-reChIP from the fraction immunoprecipitated using the anti-REST antibody confirmed TRF2 binding (Supplementary Figure 2E). ReChIP for TRF2 was negative for the *synapsin* promoter as expected. Together these confirmed TRF2-REST co-occupancy at the *hTERT* promoter. While silencing of TRF2 resulted in loss of REST occupancy (Figure 2E, F), REST silencing did not result in loss of TRF2 from the *hTERT* promoter (Supplementary Figure 2F) supporting TRF2-dependent REST recruitment. Together, these show that TRF2 binding at the *hTERT* promoter engages the PRC2-REST repressor complex, which results in H3K27me3 modification and induces a restrictive chromatin state that suppresses *hTERT* expression.

While co-IP of TRF2 with REST was clear (Supplementary Figure 2E), co-IP of EZH2 with TRF2 was not evident (Supplementary Figure 2G). As expected from earlier work we confirmed co-IP of REST with EZH2 (Supplementary Figure 2H). It is possible, therefore, that TRF2 recruits REST and EZH2 association is through REST.

### TRF2 binding on the *hTERT* promoter can be independent of telomere looping

Interaction of the chromosome 5p telomere end to the relatively close *hTERT* locus (∼1.2 Mb away) through chromatin looping was shown earlier^42^. Authors noted localization of TRF2 at the *hTERT* promoter and described it to result from physical association with telomeres. Here we sought to test more directly whether TRF2 occupancy at the *hTERT* was dependent on interaction with telomeres.

For this, we made a reporter construct by introducing the endogenous *hTERT* promoter (up to -1300 bp) upstream of the Gaussia luciferase gene. This construct was then inserted at the CCR5 safe-harbour locus ∼40 Mb away from the nearest telomere using Cas9-mediated genome editing in HEK293T cells (see Methods). Expression of Gaussia luciferase from the inserted construct was enhanced on silencing TRF2 and over expression of TRF2 DNA binding mutants (Figure 3A); as observed for the endogenous *hTERT* promoter. Next, we tested TRF2 occupancy on the *hTERT* promoter at the CCR5 locus using specific ChIP-qPCR primers designed for the inserted loci (which spanned from 113 bp upstream of TSS till 196 bp downstream; Figure 3A). TRF2 occupancy was clearly evident consistent with our observations made earlier with the endogenous *hTERT* promoter (Figure 3B). Together, these support TRF2-mediated transcriptional repression of *hTERT*.

**Figure 3.**
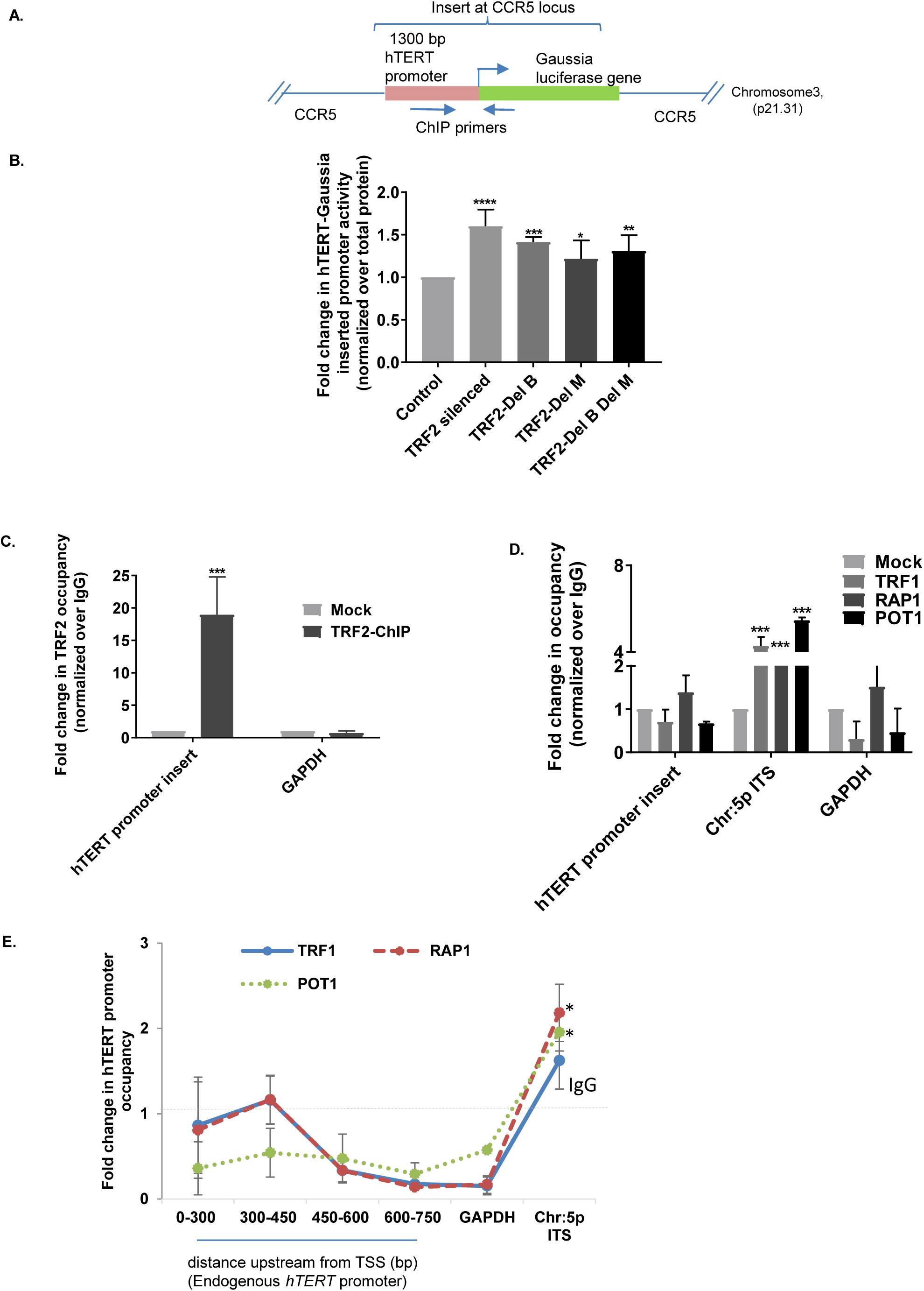
TRF2 binding on *hTERT* promoter is independent of telomere looping. **A**. Schematic for insertion of Gaussia luciferase gene, driven by (+33 to-1276) bp *hTERT* promoter inserted at *CCR5* safe harbour locus (46Mb away from nearest telomere using CRISPR/Cas9) in HEK293T cells. **B**. Effect of TRF2 silencing or over-expression of DNA binding mutants of TRF2, TRF2-DelB, TRF2-DelM and TRF2-DelB DelM on Gaussia luciferase activity, normalized over total protein **C**. qRT-PCR following TRF2 ChIP on the exogenously inserted hTERT promoter at CCR5 locus ; normalized over mock (IgG) (*GAPDH* promoter-negative control for TRF2 occupancy) **D**. qRT-PCR following ChIP for shelterin complex proteins TRF1, POT1 and RAP1, on *hTERT* promoter insert in HEK293T cells **E**. qRT-PCR following ChIP for TRF1, POT1 and RAP1 on endogenous *hTERT* promoter in HT1080 cells (in **D-E:** Chromosome 5p (interstitial telomeric sequence) ITS site-positive control and *GAPDH* promoter-negative control). All error bars represent ± standard deviations from mean values and p values have been calculated by paired /un paired T-test. (*: p<0.05, **: p<0.01, ***: p<0.005, ****: p<0.0001).

Following this we reasoned that looping/interaction with telomere ends was likely to result in presence of other telomere binding shelterin factors like POT1, TRF1 and RAP1, along with TRF2, on the inserted promoter. To test this, we selected the interstitial telomeric-like sequence (ITS) ∼100 Kb downstream of *hTERT* exon reported to engage telomeres through looping of the 5p chromosome (Kim *et al*., 2016) as positive control^43^. The GAPDH promoter was used as negative control. In contrast to TRF2 none of the other factors tested (POT1, TRF1, RAP1) showed significant occupancy at the exogenously inserted 1300 bp *hTERT* promoter (Figure 3D); whereas their occupancy was observed at the Chromosome 5p ITS sequence. Telomeric binding of POT1, TRF1, RAP1 and TRF2 was confirmed independently in each case using telomere-specific probe (Supplementary Figure 3A).

Next, we checked if TRF2 occupancy at the endogenous *hTERT* promoter was due to looping. Using the same argument as above we tested occupancy of other shelterin factors POT1, TRF1 and RAP1, which would be likely if telomeres physically associated with the *hTERT* promoter. Here also while TRF2 occupancy was clear (Figure 1A), we did not find occupancy of POT1, TRF1 or RAP1 in the region up to 750 bp upstream of the endogenous *hTERT* promoter in HT1080 cells (Figure 3E); whereas occupancy on the chromosome 5p ITS was as expected. Telomeric binding of the shelterin proteins was confirmed independently using telomere-specific qPCR probes following ChIP (Supplementary Figure 3B). Together, our results show occupancy of TRF2 at the *hTERT* promoter was independent of telomeric association and likely to be in addition to telomere-dependent effects noted earlier depending on the length of the 5p telomere.

### TRF2 binding on the *hTERT* promoter is dependent on DNA secondary structure G-quadruplex

Because the *hTERT* promoter had multiple G-quadruplexes^18–21^ (shown in Supplementary Figure 4A) we analyzed the *TERT* promoter across vertebrates. And, noted with interest that several vertebrate species have one or more G-quadruplexes within 500 bp of *TERT* TSS (Figure 4A). Previous work by us and others has reported interaction of TRF2 with G-quadruplexes from promoters as well as telomere ends^12,44,45^. Therefore, we next asked if G-quadruplexes in the *hTERT* promoter interacted with TRF2.

**Figure 4.**
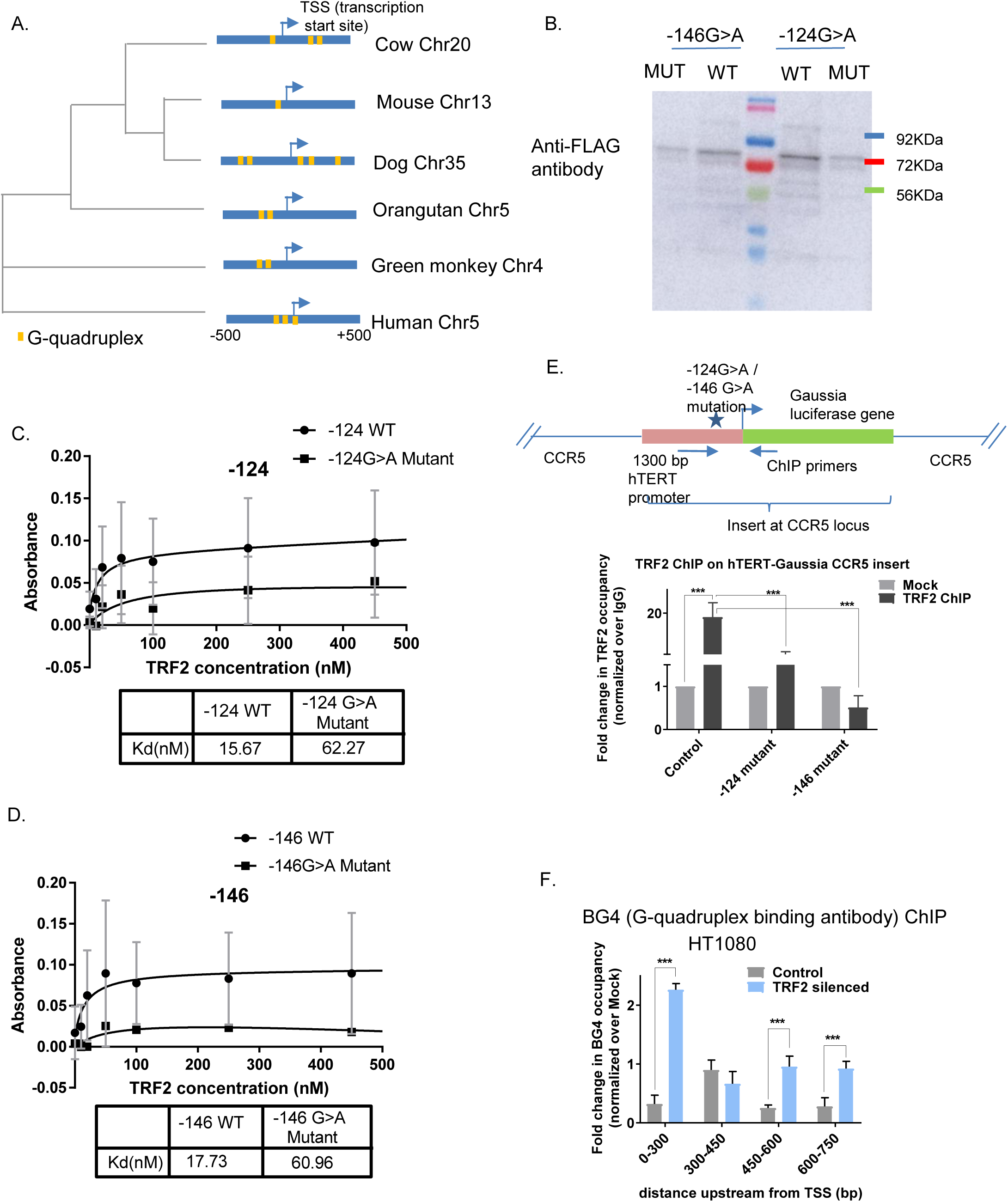
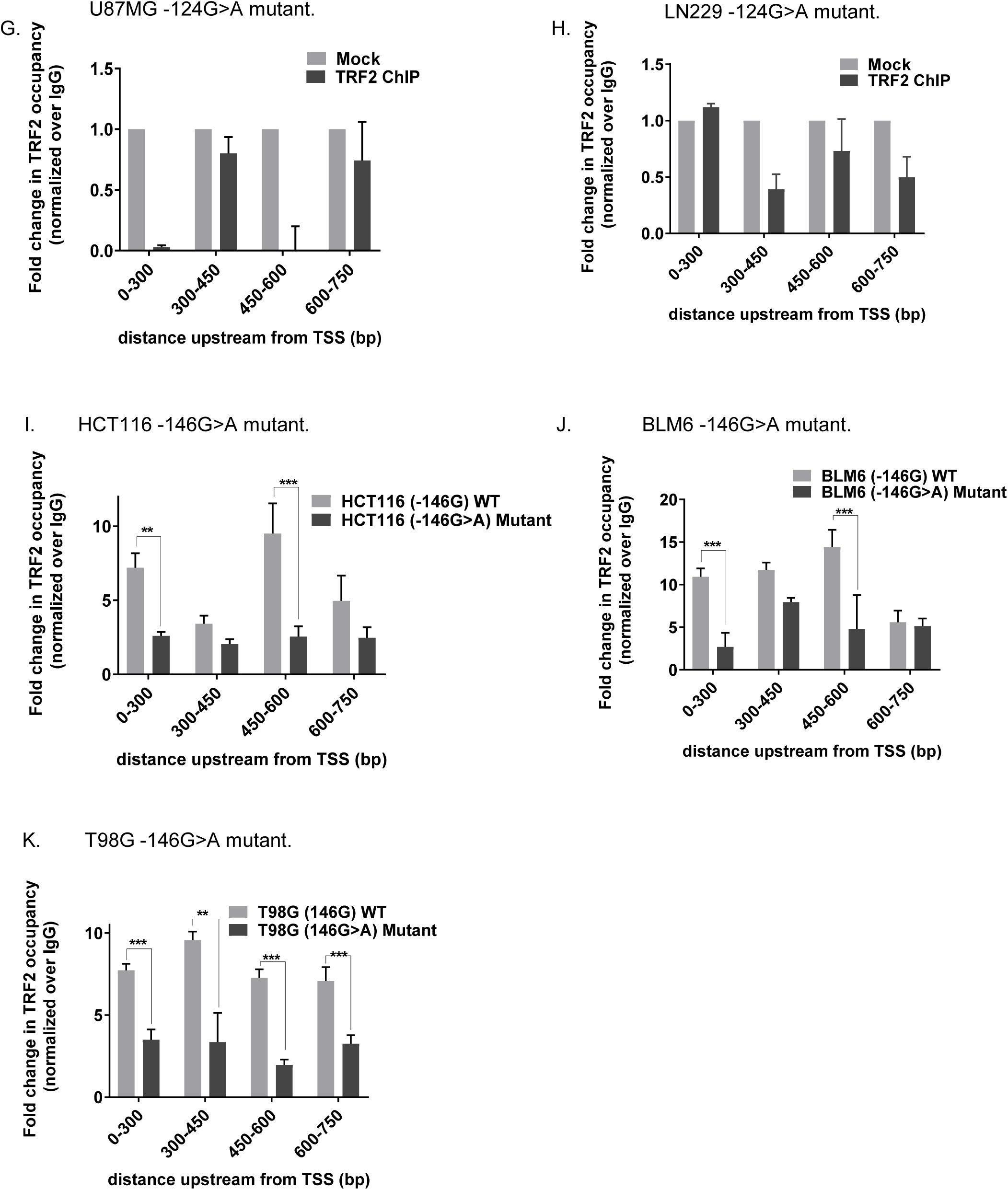
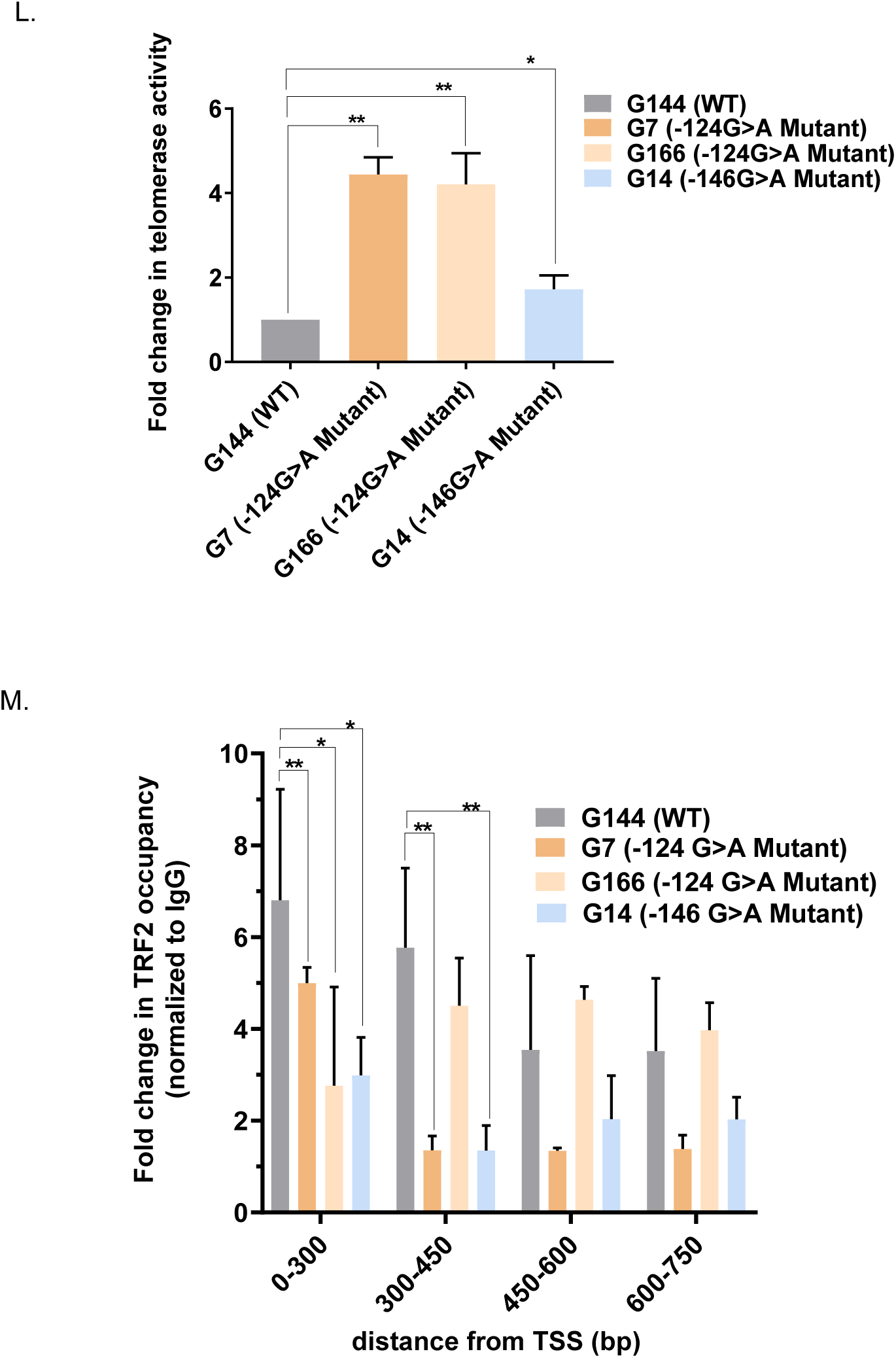
TRF2 directly binds *hTERT* promoter G-quadruplex and TRF2-mediated repression of *hTERT* is G-quadruplex-dependent. **A**. Phylogenetic tree based on the sequence spanning of ±500bp of *hTERT* TSS across vertebrates, presence of G-quadruplexes in respective organisms are shown in yellow. **B**. from whole cell lysate of Flag-tagged TRF2 over expressing HT1080 cells, oligo pull down using biotin tagged -124/146 WT and mutant oligos, followed by western blot probed with anti-flag antibody. **C-D**. ELISA experiments using increasing concentrations of purified TRF2 protein and biotin tagged – *hTERT* promoter oligonucleotides **C**. -124 WT and its G>A mutant oligo **D**. -146WT and mutant oligo. **E**. qRT-PCR following TRF2 ChIP performed on the exogenously inserted wild type and -124/-146G>A mutant *hTERT* promoter at CCR5 locus in HEK293T cells. **F**. qRT-PCR following BG4 ChIP on *hTERT* promoter spanning up-to 750bp upstream of TSS in HT1080 cells in TRF2 silenced conditions, normalized over control (both were individually normalized over respective mock). **G-H**. qRTPCR post TRF2 ChIP spanning 0-750bp upstream of endogenous *hTERT* promoter in secondary -124G>A mutant Glioblastoma cell lines, **G**. U87MG, **H**. LN229 **I-K**. q-RT-PCR following TRF2 ChIP spanning *hTERT* promoter in -146 WT in comparison to G>A promoter mutant cell lines (both were normalized over respective mock (IgG)) **I**. HCT116 WT and mutant cells **J**. BLM6 cells with WT and mutant *hTERT* promoter and **K**. T98G cells with WT and mutant *hTERT* promoter. **L**. Telomerase activity measured by ELISA TRAP and **M**. qRT-PCR post TRF2 ChIP spanning endogenous *hTERT* promoter across patient derived primary glioblastoma cells, G144 (WT *hTERT* promoter), G7,G166 (−124G>A mutant), G4 (−146G>A mutant). All error bars represent ± standard deviations from mean values and p values have been calculated by paired /un paired T-test. (*: p<0.05, **: p<0.01, ***: p<0.005, ****: p<0.0001).

First, we tested, G-quadruplex-TRF2 association at the *hTERT* promoter, we focused on two G-quadruplexes (Supplementary Figure 4A and B). Formation of G-quadruplex in solution for both the motifs was reported earlier ^21,47^. Two mutations (at -124 bp (G>A) and -146 bp (G>A) from TSS) (Supplementary Figure 4A) were found to be frequently associated with several cancers including glioblastoma (GBM) and melanomas^22,48–50^. Furthermore, both the mutations substantially destabilized the respective G-quadruplexes in solution^21^, which was further confirmed by us (Supplementary Figure 4B). To begin with, we checked TRF2 binding to the wildtype *hTERT* promoter G-quadruplexes in comparison to their respective mutants. For this, flag-tagged TRF2 was expressed in HT1080 cells. Lysate from these cells was incubated with biotinylated wild type or mutant (−124G>A or -146G>A) oligonucleotides and pulled down using streptavidin beads (Methods). Using anti-flag-antibody we observed TRF2 had enhanced interaction with wildtype relative to mutant oligonucleotides in both cases (Figure 4B). In addition, ELISA with recombinant TRF2 showed about four-fold higher affinity for *hTERT* promoter G-quadruplexes relative to the respective mutant oligonucleotides that destabilized G-quadruplex formation in both cases (Figure 4C, D).

Next, we used the *hTERT* promoter-gaussia luciferase reporter inserted at the CCR5 locus, where G>A substitutions were introduced either at the -124 or the -146^th^ positions from TSS (Figure 4E). TRF2 occupancy at the inserted *hTERT* promoter was significantly depleted in case of both the substitutions compared to the unsubstituted case (Figure 4E). As expected, TRF2 occupancy on the endogenous *hTERT* promoter remained unaltered in these cells (Supplementary Figure 4C). Taken together results suggest that for TRF2 binding at the *hTERT* promoter the two tandem G-quadruplexes tested are intact.

To further check this, intracellular presence of the *hTERT* promoter G-quadruplexes we performed ChIP using the G-quadruplex-binding antibody BG4^46^. However, we were not able to detect BG4 occupancy on the *hTERT* promoter (Figure 4F). We reasoned, as also mentioned by authors^46^, that this could be due to the presence of TRF2 on the *hTERT* promoter, which might restrict binding of BG4. Therefore, we checked for BG4 occupancy after silencing TRF2. In cells lacking TRF2 we found significant occupancy of BG4 on the *hTERT* promoter (Figure 4F). Together these support the presence of G-quadruplexes in the *hTERT* promoter, and that the G-quadruplexes might associate with TRF2 inside cells.

### TRF2 occupancy is lost in cancers with *hTERT* promoter mutations

Following the above-stated results, we tested if *hTERT* promoter G>A mutations at the -124 or the -146^th^ bp position from TSS, frequently reported to be associated with human GBM, melanoma and other cancers^22,48–50^, affected TRF2 binding. We first tested two GBM, U87MG and LN229, transformed cell lines with activated telomerase that had the -124G>A mutation in both cases^51,52^. In both cell types we could not detect any TRF2 occupancy at the *hTERT* promoter (Figure 4G, H). Further, TRF2 over expression also did not result in TRF2 binding at the *hTERT* promoter in both U87MG and LN229 cells (Supplementary Figure 4D).

Further, for the -146G>A *hTERT* promoter mutation, we tested three transformed cancer cell lines pairs with or without the mutation (gift from Tergaonkar Lab). In HCT116 colorectal carcinoma cells the -146G>A mutation was introduced resulting in telomerase hyperactivation^53^. The second and third cancer cell line pairs constituted BLM6 melanoma and T98G GBM cells where the -146G>A mutation occurred intrinsically. This was corrected by making A>G substitution in both cell lines, which gave telomerase repression as expected and reported earlier^53^.

In all the three cases, TRF2 occupancy was significantly lost from the *hTERT* promoter in case of the -146G>A mutation relative to the corresponding cell line without the mutation (Figure 4 I, J, K). We earlier noted loss/gain of H3K27me3 to be TRF2-dependent (Figure 2A, B). Here we checked this taking HCT116 cells as a candidate case – loss of H327me3 from the *hTERT* promoter in cells with the mutation (as expected from loss of TRF2) was clearly observed relative to the HCT cells that had no mutation, along with gain in H3K4me3 as reported earlier by Akincilar *et al*.^53^ (Supplementary Figure 4D).

For further confirmation, we studied primary cells from grade-four GBM patients. Upon sequencing the *hTERT* near promoter (38bp downstream to 237bp upstream of ATG) we found two cases with -124G>A mutation (G7, G166); one with -146G>A mutation (G14); and one case with no mutation (G144). Telomerase activity, as expected, was several-folds higher in GBM cells with either -124/-146 G>A mutation (G7, G166 or G14) compared to G144, which had no mutation in the *hTERT* promoter (Figure 4 L). Consistent with earlier findings we found significant loss in TRF2 occupancy at the *hTERT* promoter in G7, G166 and G14 cells relative to G144 primary GBM cells (Figure 4 M).

These show that in multiple transformed and primary patient-derived GBM/melanoma cells, TRF2 occupancy was substantially reduced in case of G>A mutations in the *hTERT* promoter. Taken along with other findings, this suggest that telomerase hyperactivation, frequently found in cancers with -124/-146 mutations in the *hTERT* promoter, causally associated with high grade GBM, melanoma and other cancers ^22,48–50^might be due to loss of TRF2-mediated repression of *hTERT*.

### Stabilization of G-quadruplex using ligands reinstatesTRF2 binding and repressor chromatin

We tested four reported intracellular G-quadruplex binding ligands^54–57^ in LN229 cells (−124G>A mutation in the *hTERT* promoter) (Supplementary Figure 5A, 5B Supplementary Table 1). Two ligands, SMH1-4.6 and JD83 showed relatively more effect on *hTERT* repression. In U87MG cells treatment with SMH1-4.6 or JD83 resulted in ∼40-50% repression of *hTERT* (Figure 5A). TRF2 expression remained relatively unaltered in presence of the ligands in both the cell lines (Supplementary Figure 5B). SMH1-4.6 and JD83 also suppressed telomerase activity in both U87MG and LN229 cells (Figure 5B).

**Figure 5.**
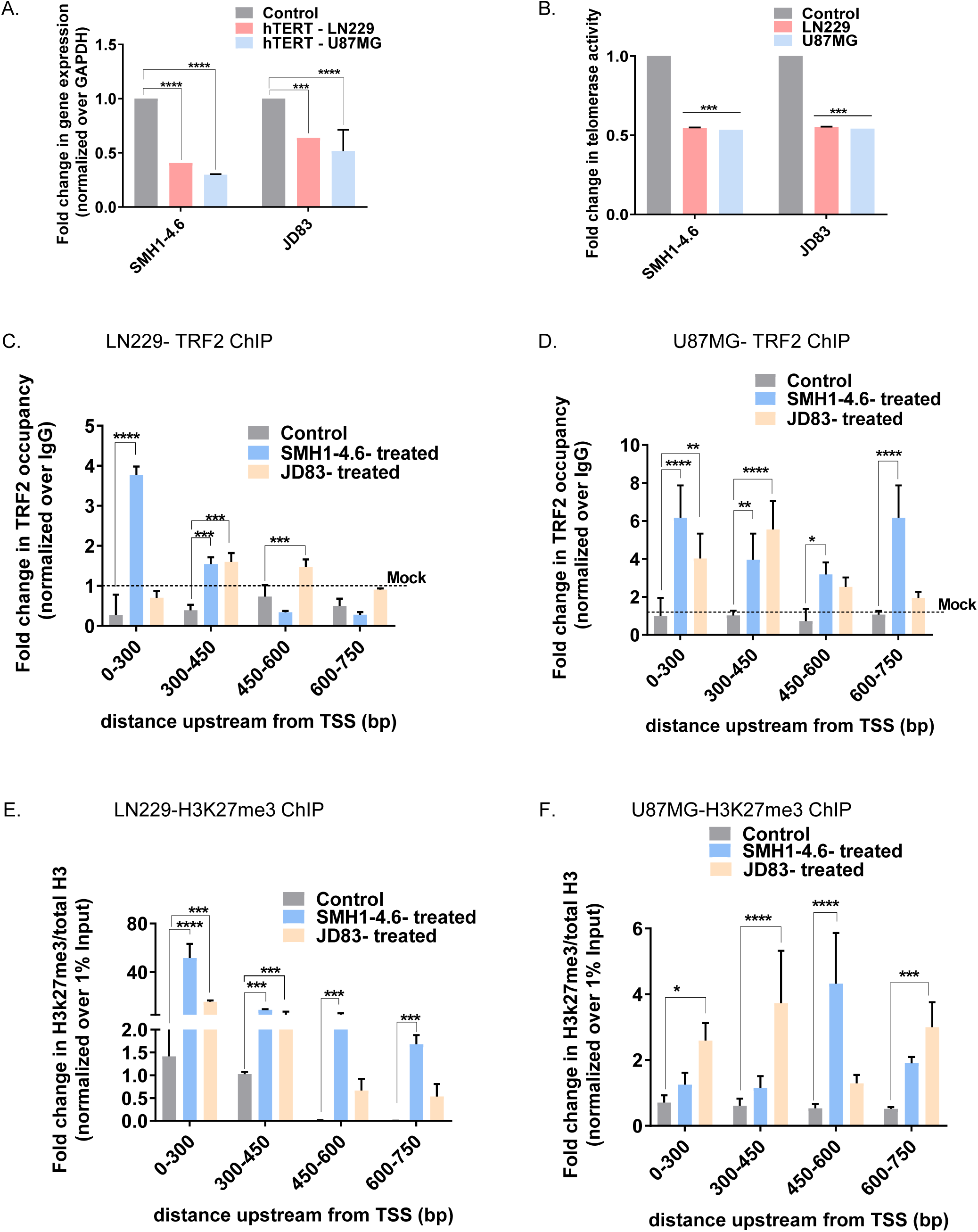
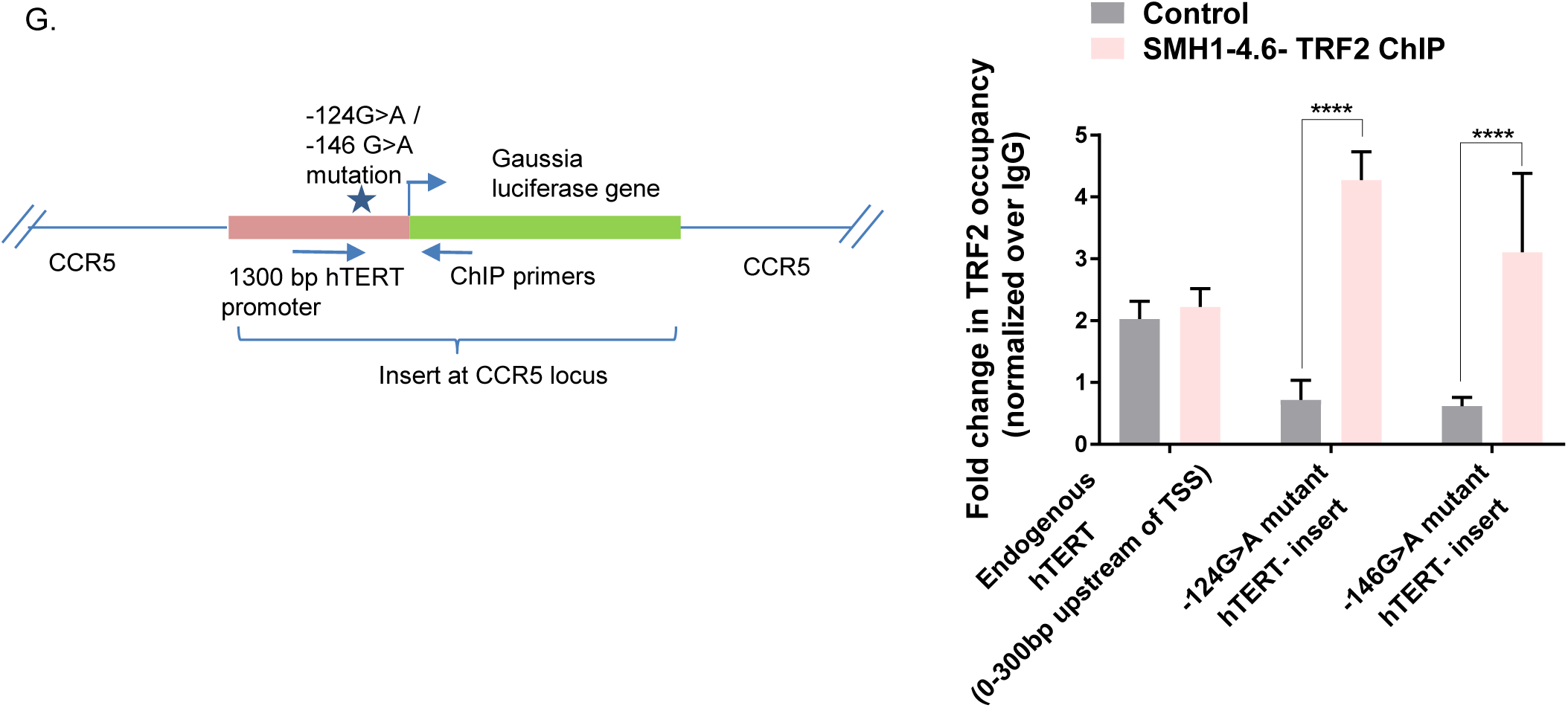
G-quadruplex binding ligands re-suppress hyperactivated *hTERT* in aggressive glioblastoma with mutations in *hTERT* promoter. **A**. *hTERT* gene expression and **B**. telomerase activity from U87MG and LN229 (−124G>A mutant) cells post treatment with SMH1-4.6 and JD83, G-quadruplex binding ligands at 2.5uM concentration for 24 hrs. **C-D**. Gain of TRF2 binding tested by qRT-PCR post TRF2 ChIP on treatment with G-quadruplex binding ligands SMH1-4.6 and JD83 ligands for 24hrs at 2.5uM concentration in -124G>A mutant cell lines **C**. LN229; **D**. U87MG cells; **E-F**. Fold change in H3K27me3/total H3 normalized over 1% Input spanning endogenous *hTERT* promoter mutant cells post ligand treatment **E**. LN229. **F**. U87MG cell lines; **G**. Gain in TRF2 occupancy on exogenously inserted *hTERT* promoter with -124 /-146G>A mutations at *CCR5* locus in HEK293T cells post treatment with SMH1-4.6 G-quadruplex binding ligand for 24hr at 2.5uM concentration, while the TRF2 occupancy remained unaltered on endogenous *hTERT* core promoter (0-300bp upstream of TSS) (*=p<0.05,**=p<0.01,***=p<0.005,****=p<0.0001).

Next, we asked whether and how treatment with the G-quadruplex-binding ligands affected TRF2 occupancy on the *hTERT* promoter. In case of both SMH1-4.6 and JD83, there was significant increase in TRF2 occupancy at the *hTERT* promoter in LN229 and U87MG cells (Figure 5C, D). Based on our results showing TRF2-mediated gain in H3K27me3 modification (Figure 2A, B), we next checked and found significant increase in the H3K27me3 histone repressor mark at the *hTERT* promoter in both LN229 and U87MG in presence of SMH1-4.6 and JD83 (Figure 5E, F). To further ascertain the effect of G-quadruplex binding ligands on TRF2 occupancy we used the *hTERT* promoter gaussia reporter (with or without -124/-146 mutation) inserted at the CCR5 locus. Treatment with SMH1-4.6 resulted in significant increase in TRF2 occupancy on the inserted hTERT promoter (Figure 5G). Taken together, this suggests ligand-mediated G-quadruplex stabilization results in recovery of TRF2 occupancy, gain in histone repressor H3K27me3 and consequent telomerase suppression in cells with *hTERT* promoter mutations that disrupt G-quadruplex formation.

## Discussion

Taken together, our results support TRF2-dependent recruitment of the PRC2-REST repressor complex maintains a non-permissive state of chromatin at the *hTERT* promoter through repressor histone modifications. This results in TRF2-mediated repression of *hTERT* expression. Experiments using recombinant TRF2 and TRF2-mutants devoid of DNA binding support transcriptional repression through direct DNA binding by TRF2 at the *hTERT* promoter. In addition, cell lines with *hTERT* promoter mutations (using CRISPR-Cas-mediated single-base editing) and an exogenous reporter cassette with specific base substitutions which disrupt the TRF2 binding site, where TRF2-mediated *hTERT* repression was lost, further confirmed transcriptional role of TRF2 in *hTERT* regulation.

TRF2 binding on the *hTERT* promoter was dependent on the presence of promoter G-quadruplex. Multiple lines of evidence reported herein support this. First, recombinant TRF2 binds with nanomolar affinity to G-quadruplexes from the *hTERT* promoter in solution, which was significantly lower when single-base substitutions are made, that disrupt the G-quadruplex stability (Figure 4C, D). Second, association of the antibody BG4, reported widely to be specific for intracellular G-quadruplexes, was found at *hTERT* promoter only when TRF2 was silenced likely because TRF2 association with the G-quadruplex competes with BG4 suggesting intracellular TRF2-G-quadruplex interaction (Figure 4F). Third, base substitutions that disrupt G-quadruplex when incorporated into the exogenously inserted promoter-reporter resulted in significantly decreased TRF2 occupancy, relative to the un-substituted promoter (Figure 4E). Fourth, in mutant cell lines with or without base substitution(s) (identical to the ones in binding assays with recombinant TRF2) that disrupt G-quadruplexes in the *hTERT* promoter, TRF2 association was significantly lower in case of cells with the G-quadruplex-disrupting substitution(s) (Figure 4 G-M). Further, this is consistent with work by us showing TRF2-G-quadruplex interactions at the *p21, PCGF3* promoters and more recently in a study where TRF2-G-quadruplex binding was found to be spread throughout genome^12,44,58^. Earlier a truncated version of TRF2 was noted to bind the telomeric G-quadruplex in solution^45^.

Based on this we sought to understand whether *hTERT* promoter mutations frequently found in aggressive GBM and melanoma that result in *hTERT* activation^22,48–50^– reported to destabilize G-quadruplexes^21^ – might disrupt interaction of TRF2 with the *hTERT* promoter. Using cancer cell lines as well as glioblastoma patient-derived primary cells harbouring G-quadruplex-disrupting *hTERT* promoter mutations we tested this: loss of TRF2 binding in case of cells with *hTERT* promoter mutation(s), but not otherwise, along with decrease in repressor histone modification (H3K27me3) and enhanced *hTERT* expression was clear in all the cases. We further tested the possibility that intracellular G-quadruplex-binding ligands might stabilize the promoter G-quadruplex (in cells with promoter mutations) and thereby reinstate TRF2 binding at the *hTERT* promoter. In presence of two different types of G-quadruplex ligands, TRF2 binding and repressor chromatin at the *hTERT* promoter, and *hTERT* repression was regained, in support of the TRF2-G-quadruplex-dependent mechanism.

Although direct transcriptional repression of *hTERT* by TRF2 has not been reported earlier, high TRF2 along with low hTERT levels was observed in CD4 T lymphocytes and a osteosarcoma-derived cell line^59,60^. Smogorzewska *et al*., on the other hand, did not find any change in *hTERT* mRNA on TRF2 induction^61^. Interestingly, in our hands, although TRF2 silencing enhanced *hTERT* expression in both cancer and normal cells, TRF2 over expression did not affect *hTERT* mRNA or protein levels significantly (data not shown). It is likely, therefore, that the chromatin in the *hTERT* promoter region is maintained in a constitutively repressed state, which is disrupted on TRF2 down regulation. This is consistent with the observation that the region upstream of *hTERT* shows presence of polycomb-mediated silencing and remains repressed in 127 human tissues (analysis based on NIH Epigenomics Roadmap) - disruption of this signature resulted in telomerase activation^62^. Furthermore, recently, Stern *et al*, observed PRC2 binding on the *hTERT* core promoter in the repressed wild type allele across cancer cells^33^. In support of these, our findings herein demonstrate TRF2 silencing results in loss of TRF2-dependent binding of the PRC2-complex at the *hTERT* promoter. This disrupts the constitutive repressed state of chromatin (and loss of repressor histone modifications) inducing *hTERT* activation.

Promoter mutation(s) in *hTERT*, on the other hand, was also reported to either generate site(s) that resulted in binding of transcription factors GABPA/B1^33,63,64^ and ETS1^65^, or RAS-ERK-mediated inhibition of HDAC1 association^66^, leading to telomerase activation in GBM, hepatocellular carcinoma and melanoma cells. GABPA along with the histone acetyl transferase BRD4 was also noted to bind the *hTERT* promoter with mutation(s) inducing permissive chromatin state^53^. Together, with the presence of tandem G-quadruplexes in the *hTERT* near promoter, the likelihood of G-quadruplex-dependent mechanisms of transcription factor association resulting in permissive/restrictive chromatin conformation has been discussed by several groups^42,43,67,68^. Consistent with this, stabilization of G-quadruplexes was noted to mask binding of the well-studied regulatory factors Sp1 and CTCF at the *hTERT* promoter^18,68,69^. From our results here, it is likely that TRF2 binding with *hTERT* promoter G-quadruplex engages the REST-PRC2 repressor complex. Loss of TRF2 occupancy in case of *hTERT* promoter mutations, on the other hand, induces open chromatin conformation allowing binding of transcription factors that re-activate telomerase expression.

Notably, *Kim et al*. observed looping of the 5p chromosome telomere to an ITS (interstitial telomere-like sequence) 1.2 Mb away and 100 kb downstream from exon1 of the *hTERT* gene^42^. This loop further engaged the *hTERT* near promoter and as a consequence telomere-bound TRF2 was physically associated to the *hTERT* promoter. Authors found loss of TRF2 binding, when telomeres were short and thereby unable to form the chromatin loop, gave increased expression of the *hTERT* transcript^42^. Our results, on the other hand, support direct binding of TRF2 to the *hTERT* promoter that is independent of telomeres (or through non-telomeric TRF2) from multiple lines of evidence. First, we reasoned that telomere-association would result in presence of other telomeric factors like TRF1, RAP1 and POT1 at the *hTERT* promoter along with TRF2 – this was not the case (Figure 5D). Second, considering the likelihood of telomere looping to diminish with physical distance we inserted an exogenous *hTERT* promoter-reporter ∼46 Mb away from the nearest telomere end – here also TRF2 association was clear whereas other telomere-bound factors were absent (Figure 5B, C). Overall this is consistent with earlier work showing non-telomeric TRF2 binding throughout the genome^12,26^. Therefore, interestingly, it is likely that both telomere-dependent^42^ and telomere-independent mechanisms of TRF2 interaction regulate *hTERT*. Further work will be required to determine in what contexts these mechanisms work, particularly in cases of ageing when telomere length changes.

In conclusion, evidence supporting TRF2-mediated re-suppression of *hTERT* using small molecule ligands in aggressive GBM and other cancers, where telomerase is hyperactivated due to *hTERT* promoter mutations, has important therapeutic potential. Perhaps more importantly, results demonstrate mechanisms of how *hTERT* is maintained in a repressed state by TRF2 in normal cells; and, that deregulation of the repression in many cases induce telomerase reactivation in cancer cells. Together, these implicate molecular connections between telomeres and telomerase, through telomeric factors like TRF2, which might be important in how normal and cancer cells manage telomeres through (de)activation of telomerase in humans.

## Materials and Methods

### Cell lines, media and culture conditions

HT1080 fibrosarcoma (ATCC-CCL-121) were maintained in Modified Eagle’s medium (MEM) supplemented with 10% Fetal Bovine Serum (FBS). HCT116 cells and TERT promoter mutant cell lines created in HCT116 background were received as a gift from Vinay Tergaonkar’s laboratory and maintained in Dulbecco’s Modified Eagle’s Medium-High Glucose (DMEM-HG) supplemented with 10% FBS. T98G Glioblastoma and BLM6 melanoma cells were also received from Vinay Tergaonkar’s laboratory and maintained in DMEM-HG with 1X Glutamax, 1X Anti-Anti (Gibco) and 10% FBS. MRC5 primary cells (ATCC-CCL171) were cultured in DMEM-HG with 1X Glutamax, 1X Anti-Anti (Gibco) and 10% FBS. U87MG (ATCC HTB-14) and LN229(ATCC-CRL-2611) glioblastoma cells were received as a gift from Dr. Ellora Sen’s laboratory and were maintained in DMEM-HG with 1X Glutamax, 1X Anti-Anti (Gibco) and 10% FBS.

HEK293T cells (ATCC-CRL-11268) and TERT CCR5 promoter insert lines made in HEK293T background were maintained in Dulbecco’s Modified Eagle’s Medium-High Glucose (DMEM-HG) supplemented with 10% FBS with addition of 1XAnti-Anti to cells post single cell seeding of CRISPR mutated pooled cell population.

### CRISPR Mediated Insertion of TERT promoter driven Gaussia Luciferase into CCR5 locus

The TERT promoter driven Gaussia Luciferase insert construct was obtained from a Genecopoeia promoter reporter clone (catalogue no- HPRM25711-PG04). The insert sequence was cloned into AY10_pS. Donor.R5.TS, a gift from Manuel Goncalves (Addgene plasmid # 100292); the donor vector sequence after cloning has been provided in supplementary information. The TERT promoter donor vector with mutation at – 124 position was generated using Quikchange SDM kit (Agilant) according to the manufacturers’ protocol. For cleavage at CCR5 locus a reported gRNA sequence (5’-GGAGAGCTTGGCTCTGTTGGGGG-3’)1 was cloned into the pX459 v2.0, a gift from Feng Zhang, that co-expresses cas9 protein and the gRNA. The gRNA cloned pX459 and the donor vector were co-transfected using FUGENE HD transfection agent according to the manufacturers’ protocol. Starting from 36 hours post transfection, the cells were treated with 2ug/ml puromycin for 3 days for selecting cells that have taken up pX459 plasmid. After growing for 5 days after selection, the cells were seeded into 96 well plates after dilution for clonal selection. 40 clones were screened using Gaussia luciferase activity and PCR to find out the positive clones (primers provided in supplementary information).

FP: TAGTGCATGTTCTTTGTGGGCT

RP: TTTTGGCAGGGCTCCGATGTAT

### Antibodies

#### Primary antibodies

TRF2 rabbit polyclonal (Novus NB110-57130), TRF2 mouse monoclonal (Millipore 4A794), TERT rabbit monoclonal (Abcam ab32020),REST rabbit polyclonal (Millipore-17-641), Histone H3 rabbit polyclonal (Abcam ab1791), Histone H3 mouse monoclonal (Abcam ab10799),H3K4me1 rabbit polyclonal (Abcam ab8895), H3K4me3 mouse monoclonal (Abcam ab1012),H3K27me3 mouse monoclonal (abcam ab6002), H3K9me3 rabbit polyclonal (Abcam ab8848),BG4 G4 specific antibody monoclonal (Millipore MABE917), EZH2 rabbit polyclonal (CST 4905),GAPDH mouse monoclonal (Santacruz 6C5).TRF1 mouse monoclonal (Novus NB110-68281), RAP1 mouse monoclonal (Santa Cruz 4C8), POT1 mouse monoclonal (Santacruz M1P1H5).

#### Secondary antibodies

anti-Rabbit-HRP(CST), anti-Mouse-HRP(CST), anti-rabbit Alexa Fluor® 488, anti-mouse Alexa Fluor® 594 (Molecular Probes, Life Technologies).

### Analysis of sequence data for detecting conserved PG4 motifs

PG4 motifs were identified from different custom fetched sequences from TERT upstream promoter region from various mammalian clades using Quadbase 2. Sequence homology and conservation scores were determined using neighbour joining cluster generation algorithm in the publicly available multiple sequence alignment tool MUSCLE.

### ChIP (Chromatin Immunoprecipitation)

ChIP assays were performed as per protocol previously reported in Mukherjee *et al*, 2019. ChIP assays were performed using relevant primary antibody. IgG was used for isotype control in all ChIP experiments. Three million cells were harvested and fixed for each were fixed with ∼1% formaldehyde for 10 min and lysed. Chromatin was sheared to an average size of ∼200-300 bp using Biorupter (Diagenode). 10% of sonicated fraction was processed as input using phenol–chloroform and ethanol precipitation. ChIP was performed using 3 μg of the respective antibody incubated overnight at 4°C. Immune complexes were collected using herring sperm DNA-saturated Magnetic Dyna-beads (protein G/A) and washed extensively using a series of low salt, high salt and LiCl Buffers. The Dynabeads were then resuspended in TE(Tris-EDTA pH 8.1) buffer and incubated with proteinase K at 55° C for 1hr. Then, phenol-chloroform-isoamyl alcohol was utilized to extract DNA from the proteinase K treated fraction. DNA was precipitated by centrifugation after incubating overnight at -20 ° C with isopropanol with glycogen and 3M sodium acetate. The precipitated pellet was washes with freshly prepared 70% ethanol and resuspended in nuclease free water. ChIP DNA was further validated by qRT-PCR method.

The primer sequences for ChIP-qPCR on TERT promoter is as follows:

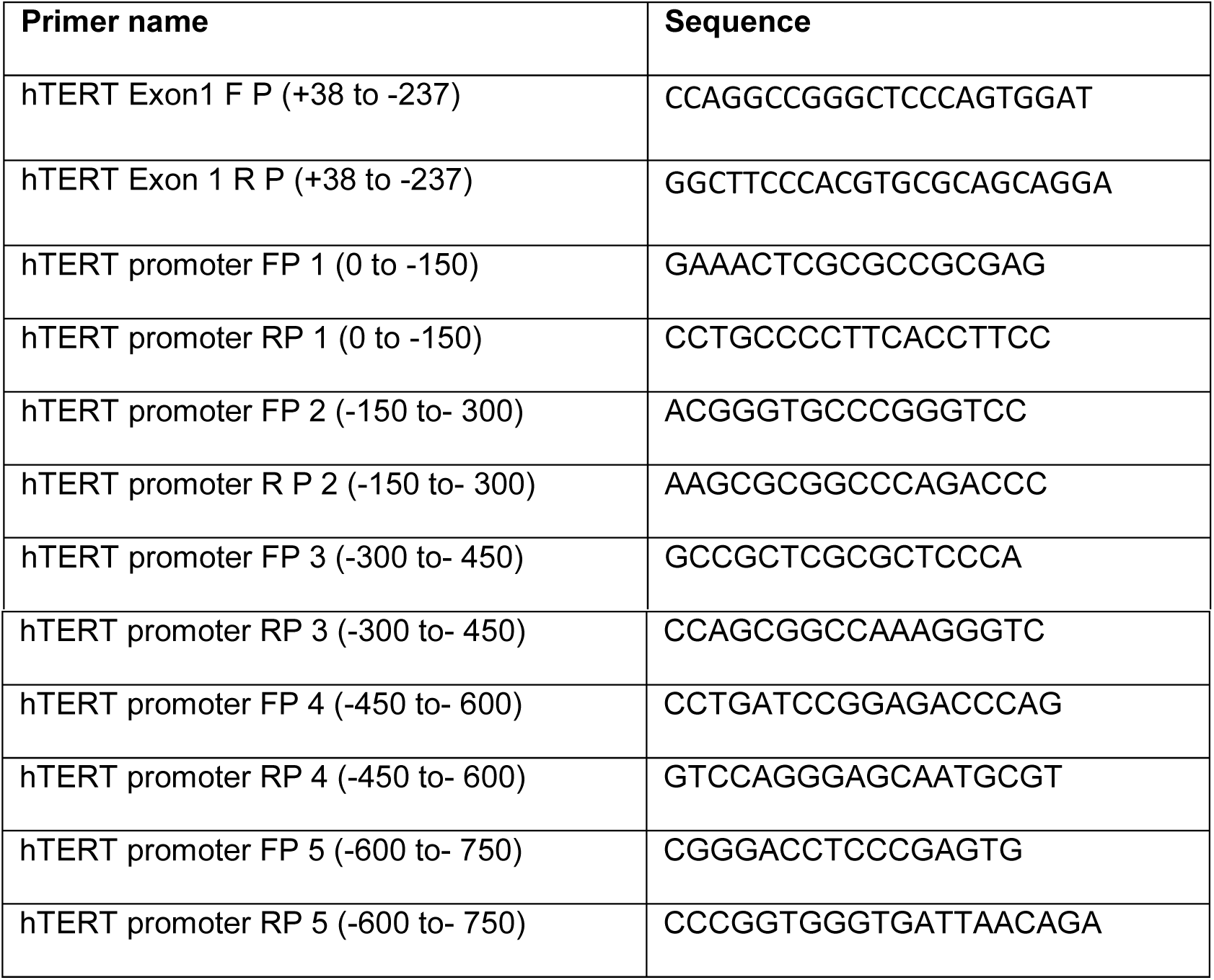

TERT ITS:

FP: GGAGCTGTGGTCTGTGTCTC RP: ACGCTAACCCTAACCCACAG

Synapsin promoter:

FP: GGTGCTGAAGCTGGCAGT RP: TGGGTTTTAGGACCAGGATG

CTCF promoter:

FP: CCTTCCAGTCGTCGCTCGCG RP: AGGAGGCTCCCGGGCCCG

GAPDH promoter:

FP: CCCAAAGTCCTCCTGTTTCA RP: GGAAGGGACTGAGATTGGC

hTERT-Gaussia ChIP primers

FP: GACCGCGCTTCCCACGTGGCGGAG RP: GCCTCGGCCACAGCGATGCAGATCAG

#### Re-ChIP

For Re-ChIP of TRF2 immunoprecipitated fraction with REST, the above stated ChIP protocol was followed with a starting harvest of 6 million cells with pull-down of TRF2 mouse monoclonal antibody using protein G dynabeads. For Re-ChIP, half the pull-down fraction was resuspended in TE buffer with 10mM DTT after the salt buffer washes and incubated for 30 mins at RT. Following this the fraction was centrifuged at 10K rpm at 4° C for 10 mins and supernatant was used as lysate for REST ChIP using REST rabbit monoclonal antibody and pull down using protein A Dynabeads.

### Immunoprecipitation of proteins

Six million cells were collected and washed in cold 1X PBS and lysed using RIPA (sigma) with 1x mammalian Protease inhibitor Cocktail as per manufacturer protocol. For immunoprecipitation experiments 1 mg of protein was incubated for 4 hours at 4°C with primary antibody in ratio recommended by manufacturer for immunoprecipitation. The pull-down was performed using Catch and Release co-immunoprecipitation kit (Millipore) as per manufacturer’s protocol.

### Immunofluorescence microscopy

Adherent cells were seeded on coverslips and allowed to reach a confluency of ∼ 70%. Cells were fixed using freshly prepared 4% Paraformaldehyde by incubating for 10 min at RT. Cells were permeabilised with 0.5% Triton™ X-100 (10 min at RT) and treated with blocking solution (5% BSA in PBS) for 2 hrs at RT. All the above stated were followed by three washes with ice cold PBS for 5 mins each. Post-blocking, cells treated with relevant antibodies as follows: anti-TRF2 antibody mouse (1:1000), anti-TERT antibody rabbit (1:1000) and incubated overnight at 4° C in a humid chamber. Post-incubation, cells were washed alternately with PBS and PBST three times and probed with secondary Ab [rabbit Alexa Fluor® 488(1:1000) / mouse Alexa Fluor® 594(1:1000)] for 2 hrs at RT. Cells were washed again alternately with PBS and PBST three times and mounted with Prolong® Gold anti-fade reagent with DAPI. Images were taken on Leica TCS-SP8 confocal microscope. LEICA LAS-AF software was used to calculate TRF2 and TERT signal intensity (a.u.).

### Immuno-Flow cytometry

3 million cells for each condition were fixed using 4% formaldehyde for 10 mins at RT followed by 3 ice cold PBS washes for 5 mins each. Cells were permeabilised using 90% Methanol (pre-chilled) for 5 mins and followed by three ice cold PBS washes for 5 mins each. Dilution of primary antibodies were made (TERT rabbit and TRF2 mouse) in 1% BSA (in PBS) in 1:250 ratio by volume. Cells were incubated with primary antibody cocktail for 2 hrs at RT. Three ice cold PBS washes to cells (10 mins each) were given and secondary antibodies-rabbit Alexa Fluor® 488(1:1000) / mouse Alexa Fluor® 594(1:1000) in 1% BSA (in PBS) were added. Cells were incubated at RT for 1hr and given three ice cold PBS washes (10 mins each). Cells were resuspended in 0.5 ml of PBS and scored for Fluorescence intensity in an Acuuri c6 flow cytometer in the FL1 (488 nm) and FL3 (594 nm) channels. The FCS files were analyzed using Flow-Jo (version 10) software.

#### Transfections

Cells were transfected using protocols previously described for TRF2 WT and mutant mammalian expression plasmids and TRF2 siRNA pool was used for TRF2 silencing 5’GGC-UGG-AGU-GCA-GAA-AUA-U3’, 5’CUG-GGC-UGC-CAU-UUC-UAA-A3’, 5’GCU-GCU-GUC-AUU-AUU-UGU-A3’ as described in Mukherjee *et al*, 2019.

### Gaussia-Luciferase assay

Minimal promoter region of TERT (∼1300 bp starting from 48 bp downstream of Transcription start site) procured from Genecopoeia-HPRM25711-PG04 (pEZX-PG04.1 vector). Gaussia luciferase kit from Promega was used for detecting secreted Gaussia luciferase signal as per manufacturer’s protocol.

#### Real time PCR for mRNA expression

Total RNA was isolated using TRIzol® Reagent (Invitrogen, Life Technologies) according to manufacturer’s instructions. RNA was quantified and used for cDNA preparation using Applied Biosciences kit. A relative transcript expression level for genes was measured by quantitative real-time PCR using a SYBR Green based method. Average fold change was calculated by difference in threshold cycles (Ct) between test and control samples. *GAPDH* gene was used as internal control for normalizing the cDNA concentration of each sample. mRNA primer sequences are as follows:

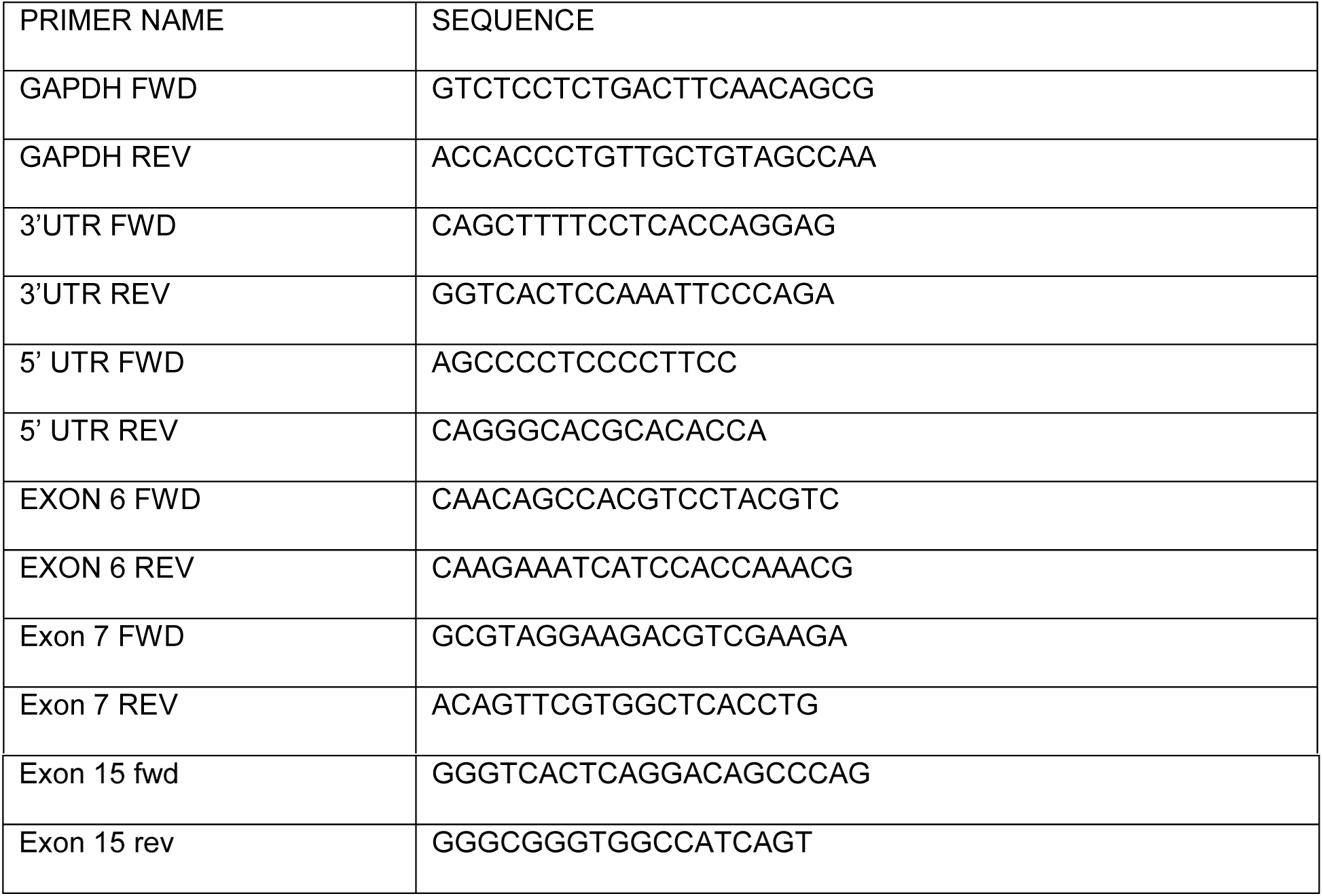

### Dot blot analysis

For dot blot analysis, Genomic/ ChIP DNA was denatured at 95°C and dot blotted on N+ hybond membrane (Amersham) in pre -wetted in 2X SSC buffer. The DNA was UV cross-linked. Membranes were pre-hybridized in Rapid-Hyb buffer (Amersham) for 30 min at 37°C. Following this, hybridization with a 24-bp radio-labeled telomeric probe (AATCCC)4 was performed for 4 hr at 37°C and membranes washed with 2X SSC and 0.2X SSC + 0.1% SDS twice at hybridization temperature before exposing overnight on phosphoimager imaging plate. All data were scanned using Bio-Rad Personal Molecular Imager. Data was processed and quantified using Image J image analysis software.

### Western blotting

For western blot analysis, protein lysates were prepared by resuspending cell pellets in passive lysis buffer/RIPA with 1x mammalian Protease inhibitor Cocktail. Protein was separated using 10% SDS-PAGE and transferred to polyvinylidene difluoride membranes (Immobilon FL, Millipore). After blocking the membrane was incubated with primary antibodies-anti-TRF2 antibody (Novus Biological), anti-TERT antibody (Abcam), anti-REST(Millipore), anti-EZH2(CST) and anti-GAPDH antibody (Santa-cruz). Secondary antibodies, anti-mouse and anti-rabbit HRP conjugates were from CST. The blot was finally developed by using Millipore HRP chemiluminescene detection kit and images in a GE chemiluminiscence imager.

### Circular dichroism

The circular dichroism (CD) spectra were recorded on a Jasco-810 Spectropolarimeter equipped with a Peltier temperature controller. Experiments were carried out using a 1mm path-length cuvette over a wave length range of 200-330 nm. 5 µM oligos were diluted in 10mM Tris HCl ph7.5 and 140mM KCl and denatured by heating to 95°C for 5 min and slowly cooled to 25°C for overnight. The CD spectra reported are representations of three averaged scans taken at 25 °C and are baseline corrected for signal contributions due to the buffer.

The oligonucleotides used for the CD experiments are as follows:

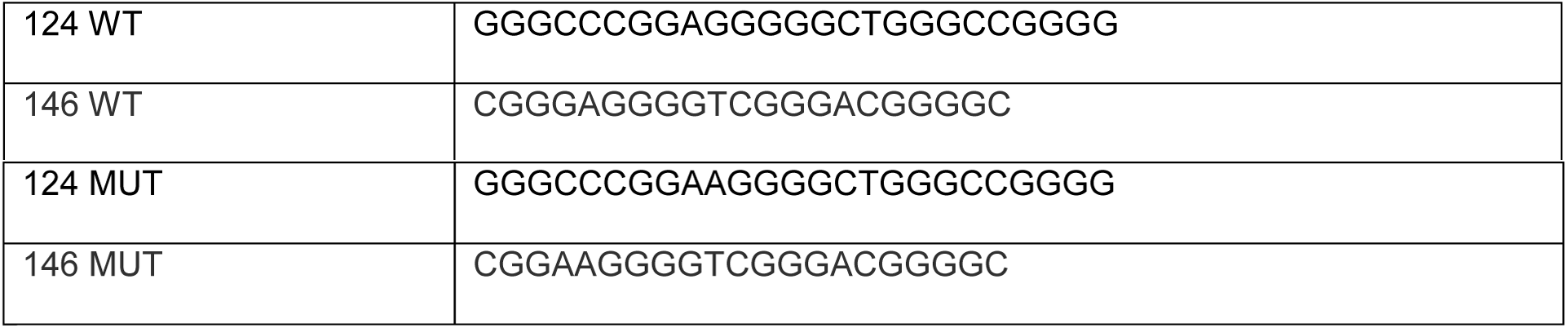

### ELISA

Biotinylated oligonucleotides were prepared at 5 µM concentration in 10mM Tris HCl ph7.5 and 140mM KCl buffer and denatured at 95 °C for 5 minutes, followed by slow cooling to room temperature to induce G-quadruplex formation. 96-well streptavidin coated pre-blocked plates from Thermo Scientific (Pierce) were used for ELISA assay. Biotinylated oligos were diluted to 5 pmol in 1X TBST buffer and loaded into each well. Oligos were incubated at 37°C on shaker for 2 hours to allow streptavidin and biotin binding and then washed 3 times with 1X TBST buffer. TRF2 protein was diluted in 1X PBST buffer and incubated with oligos for 2 hours on shaker at 4 °C and washed 3 times with 1X PBST buffer. Anti-TRF2 antibody (Novus) was used at 1:1000 dilution (60 µl per well) and incubated for 1 hr at room temperature on shaker. Wells were washed three times with 1X PBST. Alkaline phosphatase conjugated Anti-IgG antibody (Sigma) was used at 1:1000 dilution (60 µl/well) and incubated for 45 minutes at room temperature on shaker and then wells were washed once with 1X PBST and twice with 1X PBS. 30 µl BCIP/NBT substrate was added into each well and absorbance was recorded at 610 nm wavelength for 1hour with 10 min interval on TECAN multimode reader. GraphPad Prism7 was used for analysis.

#### Telomerase activity

One million cells were lysed using CHAPS lysis buffer and 1ug concentration normalized protein dilutions were used for detecting telomerase activity using ROCHE TeloTAGGG™ Telomerase PCR ELISA kit.

#### Oligo-pulldown assay

Total cell lysate of >2000ug concentration was isolated using RIPA buffer (without SDS) with 1X mPIC. Lysate was pre-cleared for cellular biotin (if any) by adding 60ul of Dynabeads MyOne Streptavidin C1 (cat no65001) beads per sample and rotating on a 4-degree celsius for 2 hours. Streptavidin beads were then removed using a magnetic stand and the lysate was divided into two equal parts. To one the wild type biotinylated oligo was added, while to the other mutant oligo was added, both amounting to 50pmoles. The lysate was incubated on rotor with oligos for 16hrs at 4-degree Celsius. Thereafter the protein and DNA were cross-linked for 15min in UV crosslinker. Thereafter 100ul of Streptavidin beads were added to each tube post twice washing of beads in 1XPBST. Beads were incubated with cross-liked lysate for 2 hours. Post this, beads were separated on magnetic stand and washed twice in 1X wash buffer (20mM Tris+10mM NaCl+ Tween 0.1%). Lastly the bound protein was eluted using Elution buffer (1MTris HCl pH6.8+10% SDS+ Bitoin 25mM). The beads were re-suspended in 50ul of elution buffer and heated at 95 degree Celsius for 5 min, the buffer was then stored in fresh tube, the process was repeated with 50ul of elution buffer. Of this total eluted protein, 60ul was run on SDS PAGE gel after adding 6X protein loading dye, as in a normal western blot protocol.

#### TRF2 ChIPseq coverage on *TERT* promoter

Sorted Alignment files (BAM) forTRF2 ChIP-seq (Mukherjee *et al*., 2019) was visualized using the publicly available software IGV for Coverage of reads on *TERT* promoter with Transcription start site (TSS) defined using transcript variant NM_198253 (RefSeq).

## Supplementary Figures

**Supplementary Figure1. A**. Reads for TRF2 ChIP seq peak on *hTERT* promoter from two biological replicates in HT1080 cells. **B**. Western blot for hTERT on TRF2 silencing, in HT1080 and MRC5 cells. **C**. FACS experiment in HT1080 and its isogenic HT-super telomerase cell line to validate hTERT antibody, as expected the antibody was able to capture close to 12-fold increase in hTERT expression in super telomerase cells. **D**. FLAG-tagged DelBDelM was expressed in HT1080 cells following that, using anti-FLAG antibody, ChIP for DelBDelM showed no significant occupancy of TRF2-DelBDelM on *hTERT* core-promoter (+38 to -237bp) as expected, whereas the occupancy of wild type TRF2 on DelBDelM over expression was lost from the *hTERT* promoter.

**Supplementary Figure 2. A-B**. hTERT promoter spanning q-RT-PCR following histone ChIP for activation marks-H3K4me3, H3K4me1 and repressor mark-H3K9me3 **A**. in HT1080 and **B**. MRC5 **C**. TRF2 occupancy spanning *hTERT* promoter in HT1080 cells on EZH2 silencing **D**. Co-immunoprecipitation of REST on TRF2 pull down from HT1080 cells. **E**. q-RT-PCR following TRF2 re-ChIP done from REST ChIP fraction in MRC5 cells (*synapsin*-positive control for REST ChIP, CTCF-negative control for TRF2 and REST ChIP. **F**. *hTERT* promoter spanning qRT-PCR post TRF2 ChIP in REST silenced background in HT1080 cells. **G**. EZH2 Co-IP with TRF2 immunoprecipitation in HT1080 cells (histone H3-positive control for TRF2 co-IP). **H**. REST co-IP with EZH2 pull down from HT1080 cells.

**Supplementary Figure 3. A-B**. Dot blot following ChIP with TRF1, TRF2, POT1 and RAP1, was probed with -telomere specific probe, showing significant enrichment of telomeric DNA over IgG **A**. in HEK293T cells with exogenously inserted *hTERT* promoter-at CCR5 locus; **B**. in HT1080 cells; quantification of data is shown in the right panel.

**Supplementary Figure 4. A**. Tandem G-quadruplexes across *hTERT* core promoter upstream of translation start site; two clinical mutations -124G>A and -146G>A, known to de-stabilize promoter G-quadruplex; **B**. CD signature of -124/146 WT and G>A mutant oligos used in the *in-vitro* experiments **C**. TRF2 binding on the endogenous *hTERT* core-promoter (+38 to-237bp) in HEK293T cells with exogenously inserted *hTERT* promoter CCR5 locus. **D**. TRF2 occupancy on endogenous *hTERT* core-promoter in U87MG and LN229 (−124G>A mutant cell lines) post TRF2 over expression. **E**. qRT-PCR spanning *hTERT* promoter following ChIP for H3K4me3 activation mark and H3K27me3 repressor mark in HCT116 WT and -146G>A mutant *hTERT* promoter cell lines.

**Supplementary Figure 5. A**. Effect of G-quadruplex stabilizing ligands on *hTERT* gene expression in LN229 cells **B**. Effect of SMH1-4.6 and JD83, G-quadruplex binding ligands on TRF2 gene expression in U87MG and LN229 (−124G>A mutant) cells. **Table1**. List of G-quadruplex stabilizing ligands screened for their repressive effect on *hTERT* gene expression in LN229 cells.

## Author Contributions

**SS**-TRF2 ChIP experiments and *hTERT* expression related experiments, *hTERT* promoter activity assays, Telomerase activity assays, Immunofluorescence microscopy, ligand treatment experiments and downstream assays, Histone ChIP, BG4 ChIP, Oligo pull-down assays, ELISA, western blots for TRF2 silencing, and Immunoprecipitation for REST and EZH2, ChIP for REST and EZH2, Re-CHIP for REST, data curation, *TERT* sequence-homology across species, figures and manuscript writing; **AKM**-Histone ChIP assays, flow-cytometry, Immunofluorescence microscopy, ChIP for Shelterin components(TRF2, TRF1,RAP1,POT1), dot blots, ChIP and Re-CHIP for REST, *TERT* expression analysis,EZH2 ChIP, ChIP-seq reanalysis, data curation, figures and manuscript writing; **SSR**-generation of *CCR5-TERT* Gaussia cell lines and characterization, **SB**-TRF2 protein purification and TRF2 ChIP assays; **SL**-TRF2 ChIP and telomerase activity in patient-derived cell lines, **DP**-TRF2 ChIP in patient-derived cell lines and telomerase activity, suggestions in manuscript writing; **MV**-Circular Dichroism experiments; **ASG**-TRF2 protein purification; **MK**-image acquisition for Immunofluorescence experiments, **SC**-conceptualization, supervision, grants for funding, figures and manuscript writing.

**Acknowledgements**

Research fellowships to SS, AKM, SSR, ASG, MK (CSIR), SB (DBT), MV (DST) is acknowledged. SL acknowledges Helse Sør-Øst, Norway and Medical Student Research Program, University of Oslo, Oslo, Norway for fellowship. Authors acknowledge Balam Singh for laboratory management, Venkatesh Prasad for suggestions on immunofluorescence experiments, all members of the SC lab for their inputs/suggestions and the IGIB HPC and Core Imaging Facility. This work was supported by research grants from Wellcome Trust/DBT India Alliance Fellowship (Grant IA/S/18/2/504021), CSIR and DBT to SC and Helse Sør-Øst, Norway and Medical Student Research Program, University of Oslo, Oslo, Norway to DP.

