## Supplementary Figures for "Human Telomerase Expression is under Direct Transcriptional Control of the Telomere-binding-factor TRF2"

Supplementary Figure 1.

A.

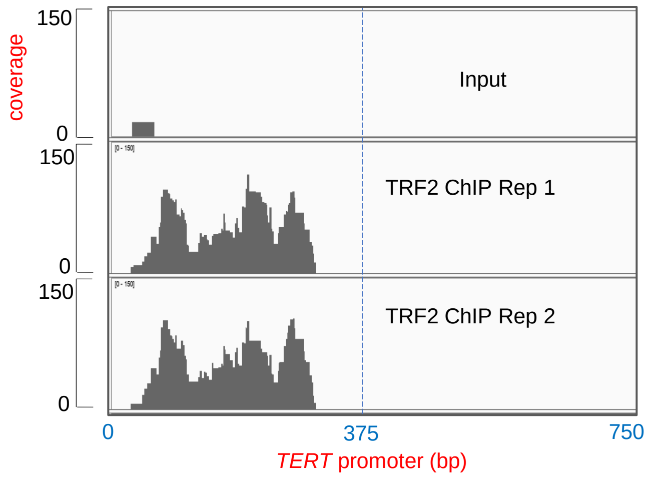


ChIP seq analysis data- TRF2 peak in *hTERT* promoter

B.

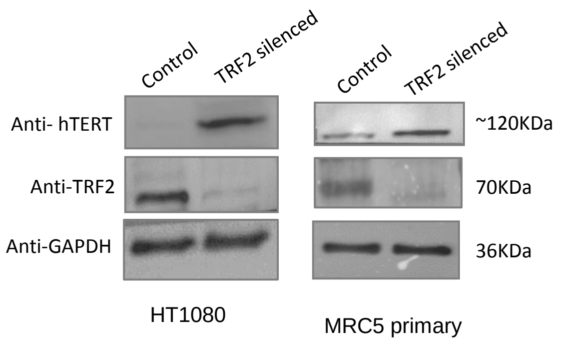


C.

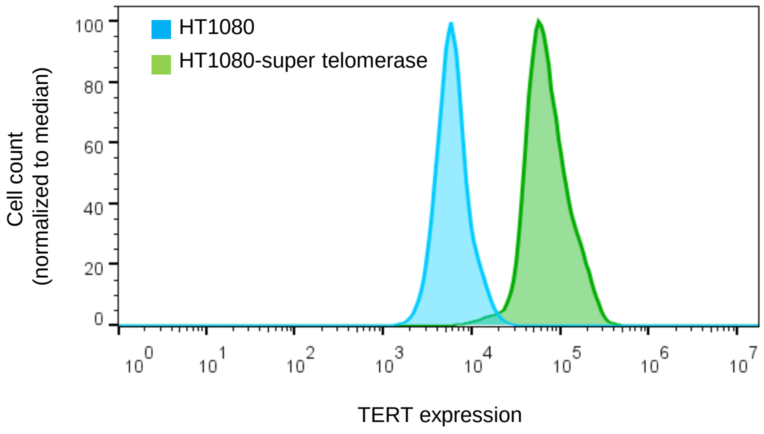


| HT1080-super telomerase | HT1080 |
|---|---|
| Mean : FL3-A: 80965 | Mean : FL3-A: 6658 |
| Robust SD : FL3-A: 40182 | Robust SD : FL3-A: 2446 |
| cells: 121806 | cells: 129816 |
| % of total cells: 89.6 | % of total cells: 92.6 |

D.

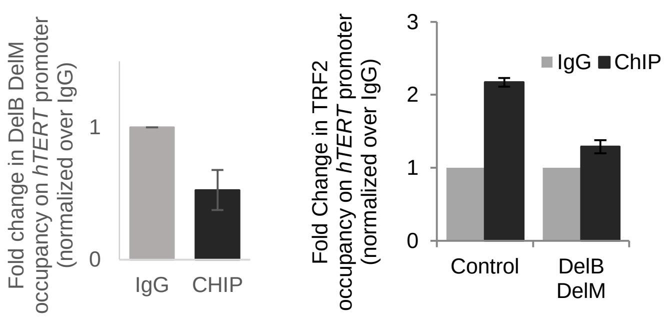

Supplementary Figure 2.

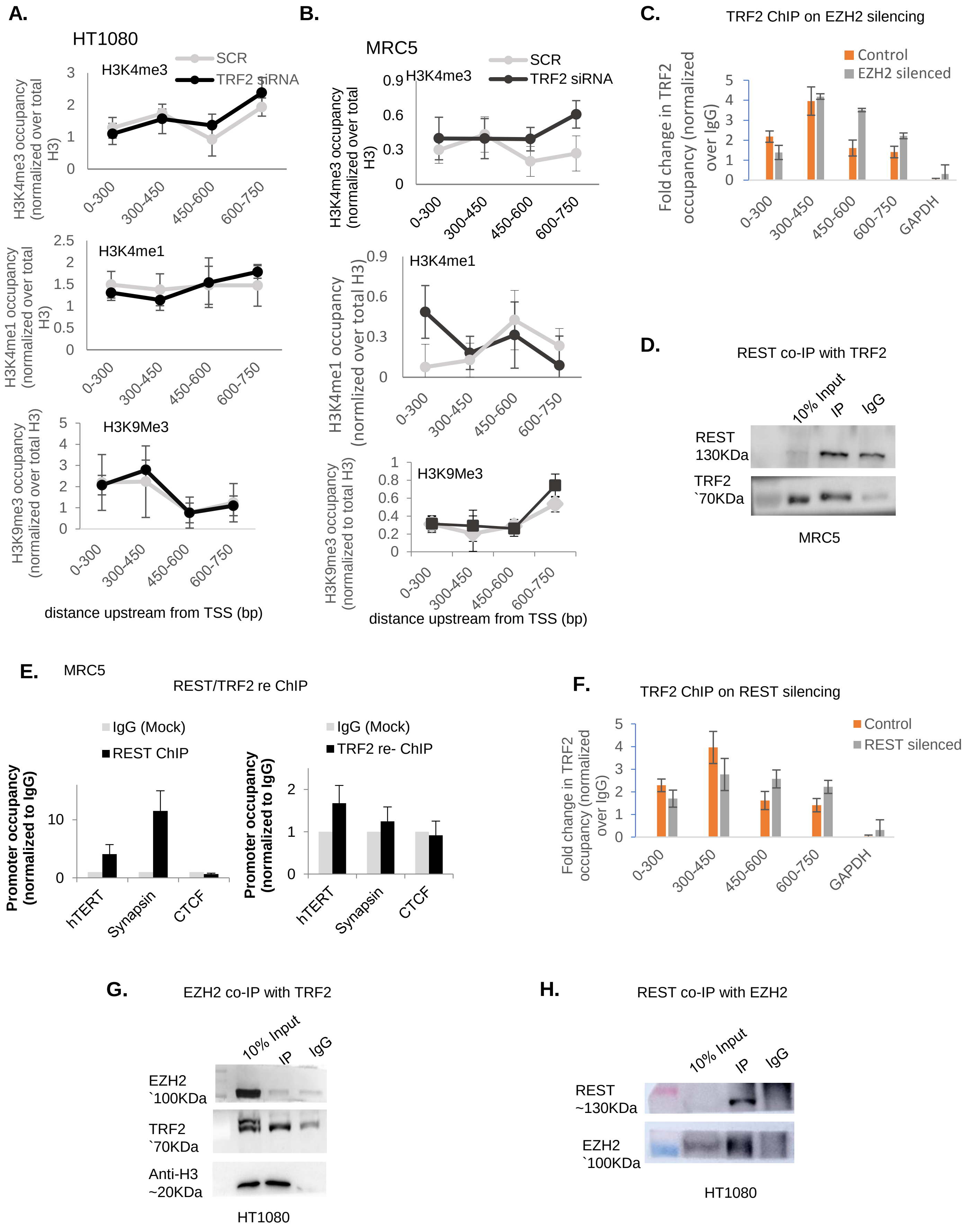

Supplementary Figure 3.

A.

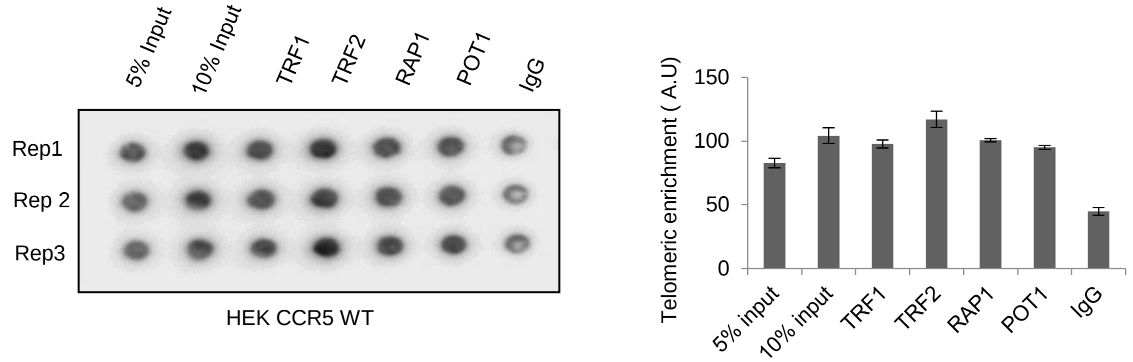


B.

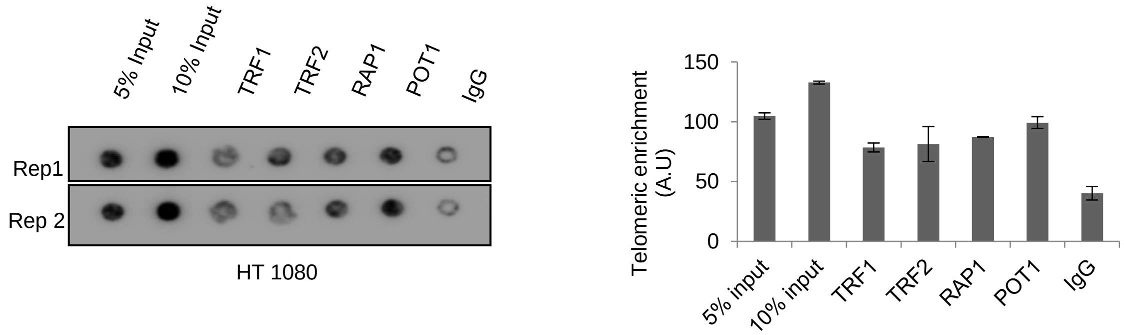

Supplementary Figure 4.

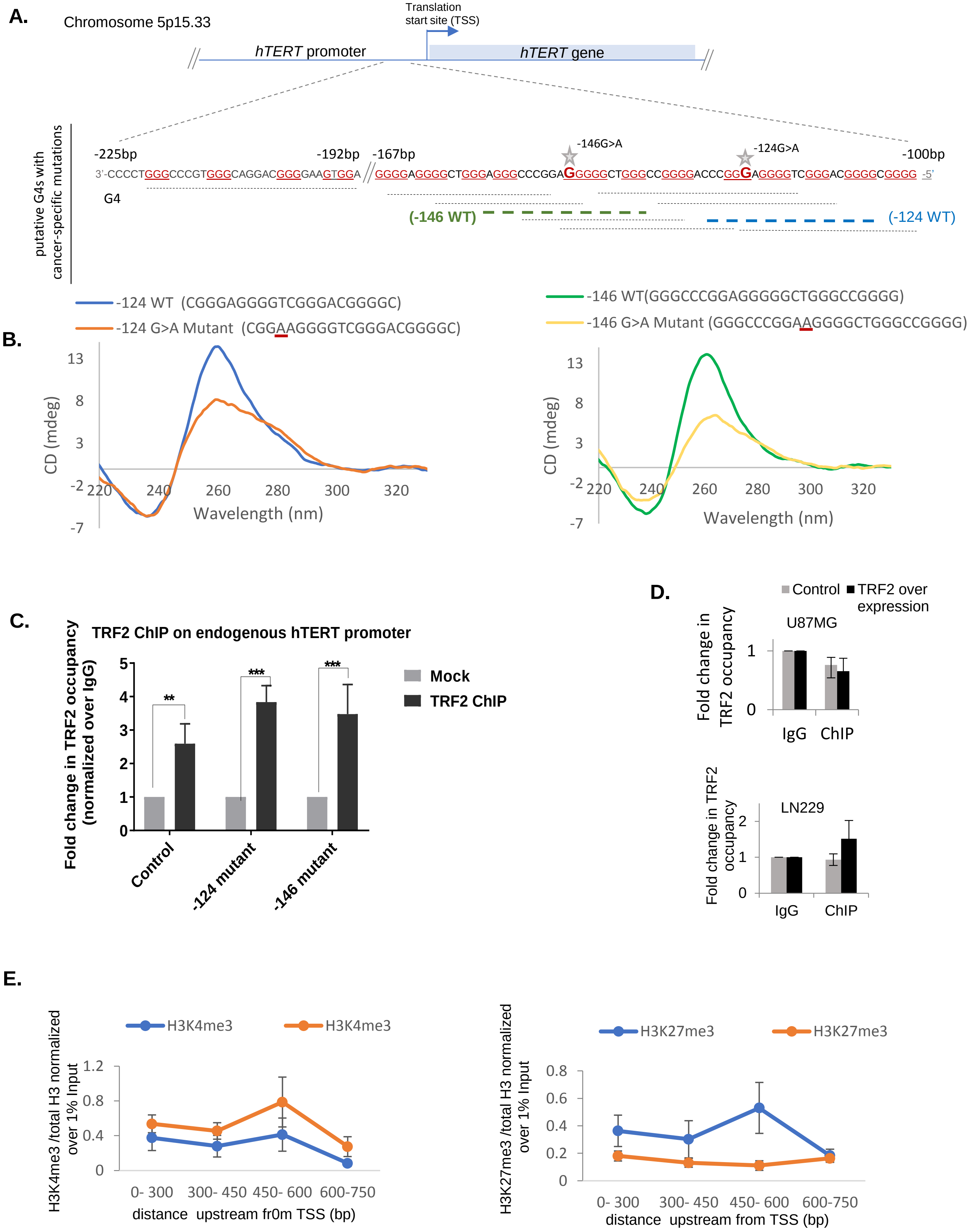

Supplementary Figure 5.

**A.**

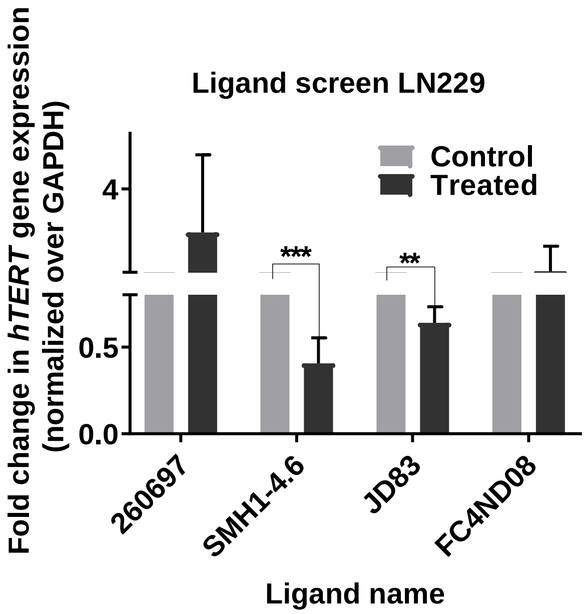


**B.**

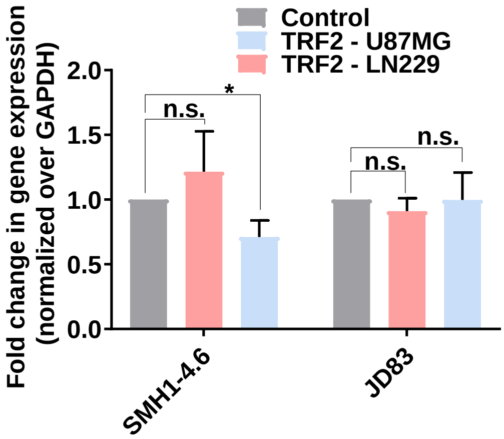


**Table 1.**

| **Ligand name** | **Reference** |
|---|---|
| 260697 | Mailliet et al 2001 patent [37] |
| FC4ND08 | Collie GW, Promontorio R, Hampel SM, Micco M, Neidle S, Parkinson GN. J Am Chem Soc. 2012 Feb 8;134(5):272331. [38] |
| SMH1-4.6 | Hampel SM, Sidibe A, Gunaratnam M, Riou JF, Neidle S. Bioorg Med Chem Lett. 2010 Nov 15;20(22):6459-63. [39] |
| JD83 | Dash J, Shirude PS, Balasubramanian S. Chem Commun (Camb). 2008b Jul 14;(26):3055-7.[40] |
